# Genome assembly, structural variants, and genetic differentiation between Lake Whitefish young species pairs (*Coregonus* sp.) with long and short reads

**DOI:** 10.1101/2022.01.15.476463

**Authors:** Claire Mérot, Kristina S R Stenløkk, Clare Venney, Martin Laporte, Michel Moser, Eric Normandeau, Mariann Árnyasi, Matthew Kent, Clément Rougeux, Jullien M. Flynn, Sigbjørn Lien, Louis Bernatchez

## Abstract

Nascent pairs of ecologically differentiated species offer an opportunity to get a better glimpse at the genetic architecture of speciation. Of particular interest is our recent ability to consider a wider range of genomic variants, not only single-nucleotide polymorphisms (SNPs), thanks to long-read sequencing technology. We can now identify structural variants (SVs) like insertions, deletions, and other rearrangements, allowing further insights into the genetic architecture of speciation and how different types of variants are involved in species differentiation. Here, we investigated genomic patterns of differentiation between sympatric species pairs (Dwarf and Normal) belonging to the Lake Whitefish (*Coregonus clupeaformis*) species complex. We assembled the first reference genomes for both *C. clupeaformis* sp. Normal and *C. clupeaformis* sp. Dwarf, annotated the transposable elements, and analysed the genomes in the light of related coregonid species. Next, we used a combination of long-read and short-read sequencing to characterize SVs and genotype them at population-scale using genome-graph approaches, showing that SVs cover five times more of the genome than SNPs. We then integrated both SNPs and SVs to investigate the genetic architecture of species differentiation in two different lakes and highlighted an excess of shared outliers of differentiation. In particular, a large fraction of SVs differentiating the two species correspond to insertions or deletions of transposable elements (TEs), suggesting that TE accumulation may represent a key component of genetic divergence between the Dwarf and Normal species. Altogether, our results suggest that SVs may play an important role in speciation and that, by combining second and third generation sequencing, we now have the ability to integrate SVs into speciation genomics.

## Introduction

Understanding the processes underlying the evolution of species and how genomes diverge during speciation is a fundamental goal of evolutionary genomics (Jiggins, 2019; Seehausen et al., 2014). The accumulation of genomic data has allowed scientists to test evolutionary scenarios and infer the timing and circumstances of species divergence (Wolf & Ellegren, 2017). Reciprocally, knowledge about the ecological, geographic, and demographic context of speciation helps to interpret the patterns of genetic differentiation between species (Jiggins, 2019; Ravinet et al., 2017). However, the genome-wide landscape of differentiation should be interpreted with caution as it results from complex interactions between gene flow, recombination, demography, and selection (Cruickshank & Hahn, 2014; Ravinet et al., 2017; Stevison & McGaugh, 2020). Analysing differentiation between evolutionarily “young” pairs of species has nevertheless proven to be informative, revealing widespread heterogeneity among and between chromosomes (Henderson & Brelsford, 2020; Martin, Davey, Salazar, & Jiggins, 2019), sometimes identifying genes underlying reproductive isolation (Hejase et al., 2020), and informing about the number and distribution of divergent loci (Dufresnes et al., 2021). Cases of ‘natural replicates’, including species pairs with similar ecological and phenotypic divergence, are of particular interest, along with instances of repeated hybridization due to secondary contacts. These instances provide important insights into the genomic architecture of species differentiation (Nadeau & Kawakami, 2019) and have revealed that similar patterns between pairs of species may be both the result of (i) shared genetic features such as low-recombination areas in which intra-specific diversity is depleted by linked selection and interspecific *F*_ST_ is inflated (Burri et al., 2015), and (ii) shared barrier loci under divergent selection or involved in reproductive isolation (Marques et al., 2016; Meier, Marques, Wagner, Excoffier, & Seehausen, 2018).

Most of our knowledge on speciation genomics is based on single-nucleotide polymorphisms (SNPs), mainly because such variants are easily accessible with short-read sequencing (Ho, Urban, & Mills, 2019; Mérot, Oomen, Tigano, & Wellenreuther, 2020). However, genomes also vary in structure with loss, gain or rearrangement of sequences between individuals and between species. Such structural variants (SVs) are now recognized to be ubiquitous and to affect a larger fraction of the genomes than SNPs (Catanach et al., 2019; Feulner et al., 2013). SVs may also have large phenotypic effects, impact recombination, and can be involved in speciation (Feulner & De-Kayne, 2017; Kirkpatrick & Barton, 2006; Wellenreuther & Bernatchez, 2018). The best recognized cases are large chromosomal rearrangements such as inversions or fusions, which are hypothesized to favour speciation by preventing recombination between alternative haplotypes (Faria & Navarro, 2010). This is supported by empirical evidence that large rearrangements can accumulate genetic incompatibilities between closely-related species of *Drosophila* (Noor et al., 2001) or fish (Berdan, Fuller, & Kozak, 2021). Whole-genome duplication events are particularly prone to favour rapid diversification (Landis et al., 2018) because the rediploidization of duplicated paralogs may differ between lineages and generate hybrid incompatibilities, as observed in yeast (Scannell, Byrne, Gordon, Wong, & Wolfe, 2006). However, small SVs, like insertions, deletions, and small duplications, may also contribute to reproductive isolation. For instance, a duplicated gene in *Drosophila melanogaster* leads to hybrid male sterility (Ting et al., 2004) while in crows a 2.25kb transposon indel underlies plumage differences, a trait involved in mate choice between two crow species (Weissensteiner et al., 2020). More generally, the insertion, deletion, duplication and/or mis-regulation of transposable elements appear to be responsible for bursts of diversification and various pre- and post-zygotic barriers, particularly in plants (Serrato-Capuchina & Matute, 2018) but also in vertebrates (Laporte et al., 2019). Overall, a better understanding of the genomic architecture of species differentiation requires the integration of SVs into speciation genomics (Feulner & De-Kayne, 2017; Mérot et al., 2020; Nadeau & Kawakami, 2019). Moreover, considering both SNPs and SVs is essential to understand the cumulative effects of those different forms of genetic variation on speciation.

Two aspects of long-read sequencing, combined with the development of new bioinformatics tools, have made it possible to investigate SVs between genomes (Ho et al., 2019; Logsdon, Vollger, & Eichler, 2020). Firstly, long-reads have improved the contiguity and quality of genome assemblies which is particularly relevant for large and complex genomes as well as for regions riddled with repeated elements (Huddleston et al., 2014). Secondly, long-reads can be directly used to detect SVs by aligning the sequences on a reference and analysing split-reads and coverage (Mahmoud et al., 2019). Together, these have proven very powerful for making catalogues of SVs within and between species. For instance, a human genome carries on average 4,442 SVs detected by short-reads (Abel et al., 2020), and 27,662 SVs detected with long-reads (Chaisson et al., 2019). Potential restrictions when generating long-reads are the requirement for high molecular weight DNA, and potentially higher costs and lower quality. Consequently, population-level analysis of SVs via long reads is not as accessible as short read sequencing. One promising possibility is to combine technologies by performing a first step of SV discovery on a limited set of high-quality samples sequenced with long-reads, and a second step of SV genotyping on more samples sequenced with short-reads (Logsdon et al., 2020; Mérot et al., 2020).

The Lake Whitefish, *Coregonus clupeaformis*, is a species complex present in numerous cold water lakes throughout North America. In the north-eastern part of the continent, it is comprised of two reproductively-isolated species, referred to as *C. clupeaformis* sp. Normal and *C. clupeaformis* sp. Dwarf, which differ ecologically by occupying the benthic and the limnetic habitat, respectively (Bernatchez et al., 2010a; Gagnaire, Pavey, Normandeau, & Bernatchez, 2013). Demographic modelling and the analysis of mitochondrial lineages showed that the two species originated from two glacial lineages that started to diverge in allopatry during the last glaciation, roughly 60,000 years ago, before coming into secondary contact about 12,000 years ago (Bernatchez & Dodson, 1990; Jacobsen et al., 2012; Rougeux, Bernatchez, & Gagnaire, 2017). This secondary contact occurred independently in several lakes of a suture zone of north-eastern America, and provoked a strong character displacement in the Dwarf species toward the use of the planktonic trophic niche, further enhancing speciation through ecological divergence (Bernatchez et al., 2010b; Landry, Vincent, & Bernatchez, 2007). The two species show limited gene flow (estimated between 1 and 30 migrants per generation in the two lakes under study (Rougeux et al., 2017)), and the rare hybrids have low fitness due to malformation, early mortality, ecological mismatch and reduced fertility (Bernatchez et al., 2010a; Renaut & Bernatchez, 2011; S. Rogers & Bernatchez, 2006). Habitat divergence is associated with species differences in a series of morphological, life-history, physiological, transcriptomic, and cytological traits (Dalziel, Laporte, Rougeux, Guderley, & Bernatchez, 2017; Dion-Côté, Symonová, Ráb, & Bernatchez, 2015; Laporte, Dalziel, Martin, & Bernatchez, 2016; Laporte et al., 2015; Rogers & Bernatchez, 2007; Rogers et al., 2002). The process of ecological and phenotypic divergence following secondary contact likely occurred independently, but with the same genetic background, in several post-glacial lakes (Rougeux et al., 2017). Multiple pairs of sympatric species thus provide valuable natural replicates to investigate parallelism and the genetic architecture of speciation. Moreover, as for all salmonid species, *C. clupeaformis* ancestors have undergone a past whole-genome duplication about 80-100 MYA followed by ongoing re-diploidization (Allendorf & Thorgaard, 1984; Lien et al., 2016; Macqueen & Johnston, 2014), resulting in a large, complex genome of approximately 2.4 to 3.5 Gb depending on the estimates (Hardie & Hebert, 2003; Lockwood, Seavey, Dillinger Jr, & Bickham, 1991). Therefore, structural genetic polymorphism is expected to be extensive in *C. clupeaformis*, though current studies have not assessed the contribution of SVs to differentiation between Dwarf and Normal species.

In this study, we used a combination of long-read and short-read sequencing (Fig. 1) to investigate the genetic architecture of speciation and address the contribution of SVs to the genomic differentiation of *C. clupeaformis* sp. Normal and *C. clupeaformis* sp. Dwarf. The main goal was to provide high-quality genomic resources for *C. clupeaformis* in order to investigate parallel and non-parallel genomic patterns of differentiation between Dwarf and Normal species in two independent North American lakes. First, we assembled the reference genome of *C. clupeaformis* sp. Normal based on one sample sequenced with long-reads and a genetic map. We documented the specificities of the genome to explore the remaining traces of previous whole-genome duplication and annotated the whitefish transposable elements. Second, we generated a catalogue of SVs varying between and within Dwarf and Normal species using three datasets: assembly comparison with a *de novo* assembly of a sympatric *C. clupeaformis* sp. Dwarf individual, high-quality long-reads of two samples (1 Dwarf and 1 Normal), and short-reads of 32 samples (17 Dwarf and 15 Normal) at medium coverage (5X). Third, we analysed genome-wide landscapes of differentiation between Dwarf and Normal species in two lakes by genotyping the whole catalogue of SVs using genome-graph based mapping, as well as SNPs, in the 32 samples sequenced with short-reads. We tested the hypothesis that the two lakes would show parallel patterns of differentiation between Dwarf and Normal and compared signals observed with different kinds of variants (SNPs vs. SVs). Our study provides a unique opportunity to characterize the contribution of both SNPs and SVs to differentiation between young species pairs, with important implications for our understanding of speciation in general.

**Figure 1:**
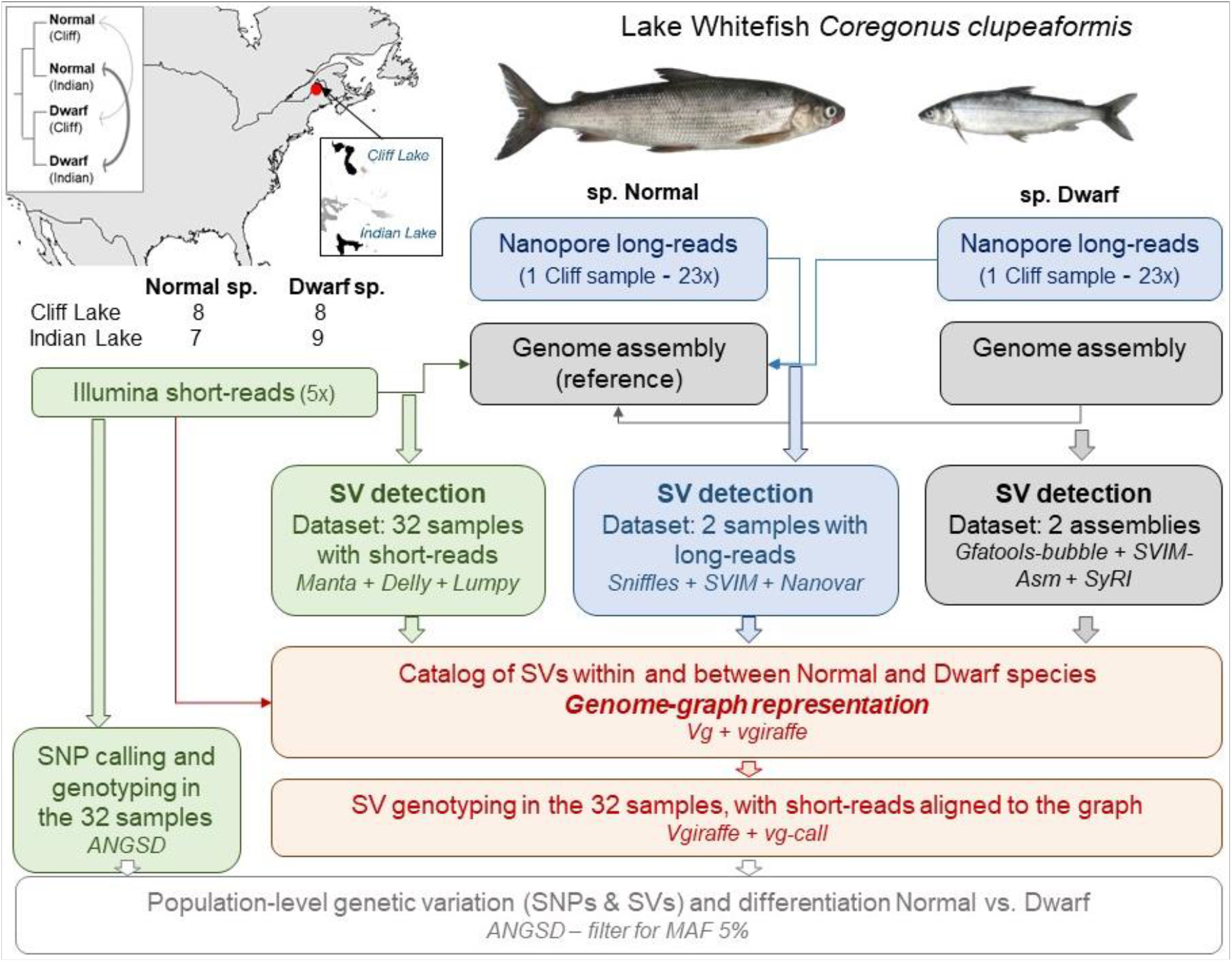
Overview of the study design. Overview of the sampling and sequencing design, which included 32 wild samples of *C. clupeaformis* Normal sp. and Dwarf sp. from Cliff Lake and Indian Lake in Maine (USA), sequenced by Illumina short reads, as well as two samples from Cliff Lake (one Normal and one Dwarf), sequenced by Nanopore long reads to assemble genomes. The insets represent the geographic locations of the two lakes sampled for this study and a schematic phylogeny of the different populations based on relationships inferred in Rougeux et al (2017), the arrows representing ongoing gene flow (1 migrant per generation in Cliff Lake, 1-30 migrants per generation in Indian Lake). The flowchart displays the main features of the pipeline of analysis performed to detect and genotype structural variants (SVs) with different datasets.

## Methods

### Sampling, DNA extraction and sequencing of *Coregonus clupeaformis*

#### Long-read sequencing

For long-read sequencing and the assembly of both reference genomes, we sampled one adult of *C. clupeaformis* sp. Normal and one adult *C. clupeaformis* sp. Dwarf from Cliff Lake, Maine (46.3991,–69.2491). Fish were caught live with gillnets, euthanized, immediately dissected to obtain fresh tissue samples, and sexed following a protocol described previously in Evans & Bernatchez (2012). Muscle samples were flash frozen in liquid nitrogen and later stored at −80 °C. High molecular weight DNA was extracted from 40mg frozen liver from both species using Qiagen Genomic Tip 100/G kit (Qiagen, Hilden, Germany). DNA integrity was assessed visually by separating fragments on a 0.5% TAE agarose gel which revealed a predominant band of high molecular weight DNA >45kb. Smaller fragments were removed by performing size selection, with >20kb cutoff, using a High Pass Plus cassette (BPLUS10) run on a Blue Pippin (Sage Scientific, MA, USA). Using 1.6ug of size selected DNA, four sequencing libraries were independently generated for each samples using the SQK-LSK109 sequencing kit (Oxford Nanopore Technologies, Oxford, UK), according to the “Genomic DNA by Ligation Nanopore” protocol. For each species, three PromethION flow cells (vR9.4.1; ONT) were loaded with library. Run performance was monitored, and once the number of sequencing pores dropped below 10% the starting number, the run was stopped and a nuclease flush was performed using the NFL_9076_v109_revA Nuclease Flush protocol from Oxford Nanopore. Additional library material was loaded onto flow-cells (by species) and sequencing initiated. In total, 3 flow cells were used to sequence the Dwarf sample (with 3 reloads among them) and 3 flow cells for the Normal sample (with 3 reloads). Raw nanopore reads were base-called using Guppy (v3.0.5. flip-flop HAC model). Data metrics before quality filtering were 72.1Gb (N50 = 27.1Kb) for the Dwarf sample and 80Gb (N50 = 27.9Kb) for the Normal sample.

#### Short-read sequencing

For population-level analysis, we sampled and sequenced 32 *C. clupeaformis* including 8 Normal and 8 Dwarf from Cliff Lake, Maine (46.3991,-69.2491), and 7 Normal and 9 Dwarf from Indian Lake, Maine (46.2574, −69.2987) with Illumina short reads. Fish were caught live with gillnets, euthanized and immediately dissected to obtain fresh tissue samples. Samples were stored in RNAlater and DNA was extracted using a modified version of a salt extraction protocol (Aljanabi & Martinez, 1997). Shotgun libraries were prepared and sequenced aiming for 5X coverage with 150bp paired-end reads on a HiSeq4000 instrument at the McGill Genome Québec Innovation center (Montréal, CQC, Canada).

Paired short reads were trimmed and filtered for quality with FastP v0.20.0 using the default parameters (Chen, Zhou, Chen, & Gu, 2018), aligned to the reference genome of the Normal *C. clupeaformis* (see below) with BWA-MEM (Li & Durbin, 2009), and filtered to keep mapping quality over 10 with Samtools v1.8 (Li et al., 2009). Duplicate reads were removed with MarkDuplicates (PicardTools v1.119.) We realigned around indels with GATK IndelRealigner (McKenna et al., 2010) and soft clipped overlapping read ends using clipOverlap in bamUtil v1.0.14 (Breese & Liu, 2013). The pipeline is available at https://github.com/enormandeau/wgs_sample_preparation

### Assembly and annotation of two reference genomes for *Coregonus clupeaformis*

#### De novo assembly and polishing

Long reads were filtered for a minimum length of 4000 bp and minimum average quality PHRED score of 7. This resulted in a total of 62.9Gb (N50=28.5kb, N90=16.3kb) for the Normal and 60.8Gb (N50=27.4kb, N90=15.0kb) for the Dwarf, hence a coverage of approximately 23x considering a genome size around 2.7Gb. For the Normal assembly, filtered long reads were independently assembled using Flye (Kolmogorov, Yuan, Lin, & Pevzner, 2019) (version 2.5, default parameters) three times using overlap sizes of 8kb, 10kb and 15kb (Table S1) . The three resulting assemblies were merged into a final assembly with Quickmerge (Chakraborty, Baldwin-Brown, Long, & Emerson, 2016) (v0.3, options: -hco 5.0 -c 1.5 -l 2000000 -ml 10000). For the Dwarf assembly, filtered long reads were assembled using Flye (version 2.5, default parameters) using overlap sizes of 8kb, 10kb and 12kb and the assembly with the best N50 was chosen (10kb). The final assemblies were first polished with their respective long reads using marginPolish (v1.2.0 https://github.com/UCSC-nanopore-cgl/MarginPolish) for the Normal and PEPPERPolish (default settings, model: pepper_r941_guppy305_human.pkl.), a successor program with similar performance, for the Dwarf. In a second step, each assembly was polished with short reads using Pilon (Walker et al., 2014) requiring a minimal coverage of 3x to polish (v1.23, --mindepth 3). BUSCO (Benchmarking Universal Single-Copy Orthologs) scores were computed to assess gene space completeness by looking for the presence or absence of highly conserved genes (BUSCO v3.0.2, reference database: actinopterygii_odb9 -sp zebrafish). BUSCO scores for the Flye polished assemblies were C:94.4%[S:50.9%,D:43.5%], F:1.7%, M:3.9%, n:4584 for the Normal and C:94.6%[S:59.1%,D:35.5%], F:0.9%, M:4.5%, n:4584 for the Dwarf. In other words, out of 4584 searched BUSCO gene groups about 94% were detected as singletons (S) or duplicates (D), a small fraction were missing (M) or fragmented (F).

#### Scaffolding into chromosomes with a genetic map

To anchor the contigs into chromosomes, we rebuilt a linkage map from previously-published data (Gagnaire, Normandeau, Pavey, & Bernatchez, 2013; S. M. Rogers, Isabel, & Bernatchez, 2007). The map is based on a backcross family whose mother is a Dwarf x Normal hybrid and father is a pure Dwarf (all details in Rogers, Isabel, & Bernatchez (2007)). The 100 full-sibs and their two parents were sequenced with reduced-representation sequencing in a previous study (Gagnaire, Normandeau, et al., 2013). Raw reads were aligned on the new contig-level assembly of the Normal genome with BWA-MEM using the default parameters (Li & Durbin, 2009). Genotype likelihoods were obtained with SAMtools mpileup (Li et al., 2009) following the pipeline and parameters provided in lep-map3 documentation (Rastas, 2017). Only positions with at least 3X coverage were kept. A linkage map was built using *Lep-MAP3* (Rastas, 2017) following a pipeline available at https://github.com/clairemerot/lepmap3_pipeline. With the *Filtering* module, markers with more than 50% of missing data, that were non-informative, and with extreme segregation distortion (χ^2^ test, *P* <10^−12^) were excluded. Markers were assigned to linkage groups (LGs) using the *SeparateChromosomes* module with increasing values of the logarithm of the odds (LOD) from 8 to 11 and a minimum size of 20 markers. Markers unassigned to LGs, or released from LG correction, were subsequently joined to LGs using the module *JoinSingle* with decreasing values of LOD until LOD=3 and a minimum LOD difference of 1. This procedure assigned 5,188 markers into 40 LGs. Within each LG, markers were ordered with 10 iterations of the *OrderMarker* module. The marker order from the run with the best likelihood was retained and refined 10 times with the *evaluateOrder* flag with 5 iterations each. To account for the lower recombination rate in male salmonids compared to females, we adjusted the parameter of recombination rates accordingly (recombination1=0.0005; recombination2=0.0025). Exploration for more stringent filtering for missing data, different values of LOD or by keeping only female-informative markers resulted in very consistent and collinear maps but with less markers, whose density is critical to accurately scaffold the genome.

Since *C. clupeaformis* sp. Normal and sp. Dwarf have the same number of chromosomes (Dion-Côté et al., 2015) and the genetic map was built from a backcross family, we used the same map to anchor both the Normal and the Dwarf genome assemblies. Scaffolds were assembled into chromosomes using *Chromonomer* (Catchen, Amores, & Bassham, 2020), which anchors and orients scaffolds based on the order of markers in the linkage map. The default parameters were used. In both assemblies, chromosomes were renamed to match homologous chromosomes in the reference genome of the European sister species, *C. lavaretus* “Balchen” (De-Kayne, Zoller, & Feulner, 2020), as detailled in Table S2. For all subsequent analyses, the Normal Whitefish genome was chosen as the reference because of its higher contiguity (N50=6.1Mb for the Normal, N50= 2.2Mb for the Dwarf) and because a higher fraction of the genome could be anchored into chromosomes in the Normal (83%) than the Dwarf (73%). It is also worth noting that, by using the same linkage map to anchor chromosomes in both the Dwarf and Normal genome, the current assemblies do not allow us to investigate large-scale chromosomal rearrangements.

#### Annotation for genes and transposable elements

Gene content annotation of both genomes was made with the NCBI Prokaryotic Genome Annotation Pipeline using the following transcriptome sources available on NCBI: Dion-Côté: PRJNA237376; Rougeux: 72 liver RNA samples from 2018, NCBI: PRJNA448004; Carruthers: SRR6321817, SRR6321818, SRR6321819, SRR6321820, SRR6321821, SRR6321822, SRR6321823, SRR6321824; Pasquier: SRP058861 Lake Whitefish, SRP045143 European Whitefish.

We used RepeatModeler2 (Flynn et al., 2020) to build a library of transposable elements from *C. clupeaformis* sp. Normal assembly. We had to slightly modify the RepeatModeler LTR pipeline because LTRHarvest failed for an unknown reason. We instead substituted it with an equivalent program, LTRfinder-parallel (Ou & Jiang, 2019), to identify LTRs in the genome. We combined the LTR-specific library with the general repeat library as done in canonical RepeatModeler2. After obtaining the TE library, we relabelled the fasta headers of sequences that were identified in the LTR pipeline but were assigned an “Unknown” classification due to lack of homology to database sequences, to broadly classify them as LTR elements.

We then used RepeatMasker to annotate the locations of each repeat family in both the Normal and the Dwarf genomes. We used parseRM.pl (https://github.com/4ureliek/Parsing-RepeatMasker-Outputs/blob/master/parseRM.pl) to summarize the genomic abundance of each TE subclass (LTR, LINE, SINE, DNA-TIR, Helitron), correcting for overlapping masking which sometimes occurs with RepeatMasker. We also used parseRM.pl to produce a landscape plot of the genome composition, where the TE-subclass composition is shown in one percent divergence windows (compared to each TE copy’s respective consensus sequence), where low divergence sequences suggest more recent insertions and higher divergence sequences suggest older insertions.

#### Synteny, map, chromosomes, and genome analysis

To analyse synteny with related species, we first compared the linkage map to the previously-published maps of *C. clupeaformis* (Gagnaire, Normandeau, et al., 2013), *C. lavaretus* “Albock” (De-Kayne & Feulner, 2018) and *C. artedii* (Blumstein et al., 2020) using MapComp (Sutherland et al., 2016), a program designed to compare syntenic relationships among markers between linkage maps of any related species using an intermediate genome, here, our reference genome. Correspondence between chromosomes and linkage groups across maps of different *Coregonus sp.* is provided in Table S2 and Fig. S1-S3.

Next, we aligned the repeat-masked *C. clupeaformis* sp. Normal and sp. Dwarf genomes to the European Whitefish reference, *C. lavaretus* sp. Balchen (De-Kayne et al., 2020), and to each other, with nucmer (-l 100 -c 500, (Marçais et al., 2018)) and used Symap v4.2 (Soderlund, Bomhoff, & Nelson, 2011) to extract syntenic blocks along the genome. Syntenic blocks were visualised in R using the package *Circlize* (Gu, Gu, Eils, Schlesner, & Brors, 2014).

To investigate chromosome types (acrocentric, metacentric), we used phased information from the linkage map by applying a method developed by (Limborg, McKinney, Seeb, & Seeb, 2016), which uses phased progeny genotypes to detect individual recombination events. The cumulative number of recombination events between the first marker and increasingly distant markers was computed from both extremities of each chromosome and this recombination frequency (RFm) is expected to reach a plateau over a chromosome arm (see Limborg et al. (2016) for details and Fig. S4).

As salmonids have experienced an ancestral whole-genome duplication, most of the chromosomes are expected to be homologous to another one, and some pairs still recombine to a certain extent, resulting in pseudo-tetrasomal regions or chromosomes (Glasauer & Neuhauss, 2014; Lien et al., 2016; Sutherland et al., 2016). To investigate this homology, we explored self-synteny by aligning the repeat-masked *C. clupeaformis* sp. Normal genome on itself with nucmer (-maxmatch -l 100 -c 500, (Marçais et al., 2018)) and extracted syntenic blocks with Symap v4.2 (Soderlund et al., 2011). The degree of sequence similarity within each of the syntenic blocks was calculated after a subsequent alignment with Lastz (Harris, 2007), following the procedure described in Lien et al. (2016). To assign *C. clupeaformis* chromosomes to ancestral chromosomes following the nomenclature proposed by Sutherland et al. (2016) based on Northern Pike (*Esox Lucius*) linkage groups, we aligned the repeat-masked Normal genome to the Northern Pike reference genome with Minimap2 (Li, 2018) and visualised alignment using D-Genies (Cabanettes & Klopp, 2018).

We further explored whether the assembly included duplicated or collapsed regions by quantifying variation of coverage along the genome. Total depth of aligned short reads across the 32 samples was calculated using ANGSD (Korneliussen, Albrechtsen, & Nielsen, 2014) at each position with the option

–doDepth –dumpCounts, and averaged by sliding windows of 100kb. The coordinates of putatively-collapsed regions, defined as regions having a depth higher than the average depth plus twice the standard deviation and showing no homology with another chromosome, is provided in Table S3.

### Detection and characterisation of SVs

We performed SV detection based on three datasets, (i) the genome assemblies of the Normal and the Dwarf, (ii) the long reads of the two samples (Normal and Dwarf) used to build the genome assemblies, and (iii) the short reads of 32 samples (Normal and Dwarf). SV detection with the three datasets shared consistent features. First, all SVs were defined relative to the reference genome of *C. clupeaformis* sp Normal. Second, to enhance SV detection, SVs were detected by three independent software packages, but to better limit the amount of false positives, we kept only SVs detected by at least 2 out of 3 SV callers in each dataset as proposed by (De Coster et al., 2019; Weissensteiner et al., 2020). Third, we focused on variants over 50bp (Ho et al., 2019) and restricted our analysis to insertions (INS), deletions (DEL), duplications (DUP) and inversions (INV) to simplify the use of multiple tools, including merging software and genome-graph representations. Fourth, to avoid artefacts due to genome misassemblies, we filtered out SVs which overlapped a scaffold junction (characterized by a gap of 10 Ns)

#### SV detection based on the comparison of de novo assemblies

SVs between the Normal and the Dwarf haploid assemblies were identified using three independent approaches detailed below. All methods included an alignment step of the query assembly (*C. clupeaformis* sp. Dwarf) on the reference assembly (*C. clupeaformis* sp. Normal). To avoid artefacts due to scaffolding with a map, we chose to use the contig-level assembly for the Dwarf genome.

i. We built a genome-graph with the two assemblies using Minigraph (Li, Feng, & Chu, 2020) with the -xggs options and retrieved SVs in a bed format with gfatools-bubble. The graph with variants was further reformatted into a vcf with full sequence information using vg suite (Hickey et al., 2020).
ii. We aligned the assemblies with Minimap2 (Li, 2018) and parameters -a -x asm5 --cs -r2k, and extracted SVs with Svim-Asm (Heller & Vingron, 2020) and the following parameters: --haploid--min_sv_size 50 --max_sv_size 200000 --tandem_duplications_as_insertions -- interspersed_duplications_as_insertions.
iii. We ordered the scaffolds of the Dwarf assembly according to the Normal reference using Ragtag (Alonge et al., 2019) and aligned the assemblies with Minimap2 (Li, 2018) and parameters “-ax asm5” and ran SyRI (Goel, Sun, Jiao, & Schneeberger, 2019) with standard parameters.

After filtering, the three VCFs were joined using Jasmine (Kirsche et al., 2021) using the following parameters “--ignore_strand --mutual_distance --max_dist_linear=0.5 --min_dist=100”, and we kept SVs detected by at least 2 approaches. All scripts are available at https://github.com/clairemerot/assembly_SV

#### SV detection based on long reads

We mapped long reads from both the Dwarf and the Normal samples to the Normal reference using Winnowmap2 v2.0 with the “--MD” flag to better resolve repetitive regions of the genome (Jain et al., 2020). SAM files were sorted and converted into BAM files with samtools V1.3.1 (Li et al., 2009). The SV detection was performed with three long-read specific SV calling programs: Sniffles v1.0.12 (Sedlazeck et al., 2018) (-l 50 -s 7 -n -1), SVIM v1.2.0 (Heller & Vingron, 2019) (--insertion_sequences) and NanoVar v1.3.9 (Tham et al., 2020) with default settings. VCF files were filtered using custom R scripts to remove excess information and read names were added to preserve insertion sequences in the final VCF. We kept SVs detected by at least two callers after merging with Jasmine v1.1.0 (Kirsche et al., 2021) including refinement of insertion sequences with Iris “max_dist_linear=0.1 min_dist=50 --default_zero_genotype --mutual_distance min_support=2 --output_genotypes --normalize_type -- run_iris iris_args=--keep_long_variants”. All scripts are available at https://github.com/kristinastenlokk/long_read_SV

#### SV detection based on short reads

SVs among the 32 samples sequenced with short reads were identified using three independent approaches: (i) Manta (Chen et al., 2016), (ii) the Smoove pipeline (https://github.com/brentp/smoove) which is based on Lumpy (Layer, Chiang, Quinlan, & Hall, 2014) and (iii) Delly (Rausch et al., 2012). All of the approaches rely on the filtered bam files resulting from the alignment of the short reads to the Normal reference (as described above). All SV callers were run with defaults parameters except for Smoove which was run by subsets of chromosomes, and Delly by subsets of individuals. VCF outputs were formatted and filtered with custom scripts called “delly_filter”, “manta_filter”, “smoove_filter” to include full sequence information. The three VCFs were joined using Jasmine (Kirsche et al., 2021) and the following parameters “--ignore_strand -- mutual_distance --max_dist_linear=0.5 --min_dist=50 --max_dist=5000 --allow_intrasample”, and we kept SVs detected by at least 2 approaches. All scripts are available at https://github.com/clairemerot/SR_SV

#### Analysis and annotation of SVs

SVs detected by the three kinds of datasets (assembly comparison, long reads, short reads) were joined using Jasmine (Kirsche et al., 2021) and the following parameters --ignore_strand --mutual_distance --max_dist_linear=0.5 --min_dist=100 --min_overlap 0.5”. This merging tool represents the set of all SVs as a network, and uses a modified minimum spanning forest algorithm to determine the best way of merging the variants based on position information (chromosome, start, end, length) and their type (DEL, INS, DUP, INV), requiring a minimum overlap between SVs and a maximum distance between breakpoints. We explored different parameters values without noticing major differences in the final merging, hence the final choice of intermediate parameters (50% of the length). We reported the overlap of SVs detected in more than one dataset according to its type and its size. The sequences included in SVs (e.g. the reference sequence in the case of a deletion, or the alternative sequence in the case of an insertion) were annotated for transposable elements using RepeatMasker and the TE library of the Normal *C. clupeaformis* (see above). We explored the length of SV sequences covered by TE or simple repeats quantitatively (Table S4-S5) and also categorized them as associated with TE or other kinds of repeats if more than 50% of the SV sequence was covered by a given TE family or other kind of repeats.

### Analysis of single-nucleotide and structural polymorphisms

#### SNPs calling and genotyping

To detect SNPs and genotype them, we analysed the short reads of the 32 samples, in bam format, with the program ANGSD v0.931 (Korneliussen et al., 2014), which accounts for genotype uncertainty and is appropriate for medium coverage whole genome sequencing (Lou, Jacobs, Wilder, & Therkildsen, 2020). Input reads were filtered to remove low-quality reads and to keep mapping quality above 30 and base quality above 20. Genotype likelihoods were estimated with the GATK method (-GL 2). The major allele was the most frequent allele (-doMajorMinor 1). We filtered to keep positions covered by at least one read in at least 50% of individuals, with a total coverage below 800 (25 times the number of individuals) to avoid including repeated regions in the analysis. From this list of variant and invariant positions, we selected SNPs outside SVs and with a minor allele frequency (MAF) above 5%. We subsequently used this SNP list with their respective major and minor alleles for most analyses, including PCA, *F*_ST_, and allelic frequency difference (AFD).

#### SVs genotyping

To genotype the identified SVs in the 32 samples, we used a genome-graph approach with the vg suite of tools (Garrison et al., 2018; Hickey et al., 2020). Briefly, the full catalog of SVs discovered (through assembly comparison and long reads and short reads SV calling) was combined with the reference genome to build a variant-aware graph using the module *vg autoindex –giraffe*. Short reads from the 32 samples were then aligned to the graph with the module *vg giraffe* (Sirén et al., 2020). For each SV represented in the graph through a reference and an alternative path, a genotype likelihood was calculated with the module *vg call*. We then combined the VCFs of SV genotype likelihoods across the 32 samples. For population-level analysis, mirroring the filters applied for SNPs, we retained SVs covered by at least one read in at least 50% of samples, and with an alternative allele frequency between 5% and 95%. The pipeline used is available here https://github.com/clairemerot/genotyping_SV. Subsequent analytical steps were performed in ANGSD, using the VCF of SV genotype likelihoods as input, to perform population-level analysis within a probabilistic framework to account for the uncertainty linked to medium coverage.

#### Genetic differentiation according to lake and species

An individual covariance matrix was extracted from the genotype likelihoods of SNPs and SVs in beagle format using PCAngsd (Meisner & Albrechtsen, 2018). The matrix was decomposed into PCs with R using a scaling 2 transformation which adds an eigenvalue correction (Legendre & Legendre, 2012). Pairwise *F*_ST_ differentiation between all populations was estimated based on the allele frequency spectrum per population (-doSaf) and using the realSFS function in ANGSD. Minor allelic frequencies per population (MAF) were estimated based on genotype likelihoods using the function doMaf in ANGSD. We then computed allelic frequency difference (AFD) between sympatric species in each lake for each variant as AFD=MAF(Dwarf) – MAF(Normal). AFD is a polarised difference of frequency that varies between −1 and 1, meaning that when we compared AFD between lakes they can be either with same sign (the same allele has a higher frequency in the same species in both lakes) or opposite sign (the allele more frequent in the Dwarf in one lake is more frequent in the Normal in the other lake). For *F*_ST_ and AFD estimates, positions were restricted to the polymorphic SNPS/SVs (>5% MAF) previously assigned as major or minor allele (options –sites and –doMajorMinor 3), and which were covered in at least 50% of the samples in each population. Given the high density of SNPs, *F*_ST_ and mean absolute AFD were also calculated by windows of 100kb for visualisation and correlation tests. The most differentiated variants between species were defined in each lake as the ones within the upper 95% quantile for *F*_ST_, and either below the 2.5% or above the 97.5% quantile for AFD. By chance only, we would expect that 0.25% of variants are in the upper *F*_ST_ quantile in both lakes (5% × 5%), 0.125% of variants are in AFD outliers in both lakes with same sign (2.5% × 2.5% × 2), and 0.125% of variants are in AFD outliers in both lakes with opposite sign. We used Fisher’s exact test to determine whether the number of outlier variants overlapping between lakes exceeded this expectation.

Using BEDtools, we extracted the list of genes overlapping with the most differentiated SNPs/SVs. We then tested for the presence of overrepresented GO terms using GOAtools (v0.6.1, *P*_val_ = 0.05) and filtered the outputs of GOAtools to keep only GO terms for biological processes with an FDR value equal to or below 0.1.

Using our annotation of TEs and repeated sequences on SVs, we tested whether some families of TEs were over-represented in the subset of outlier SVs relative to the whole pool of SVs studied at the population level using a Fisher exact test.

Finally, several Quantitative trait loci (QTLs) for behavioural, morphological and life-history traits differentiating Normal and Dwarf previously identified in Gagnaire, Normandeau, et al. (2013) and Rogers et al. (2007) were positioned on the Normal reference genome. We compared the positions of those QTLs relative to the most differentiated regions and extracted the list of genes hit by an outlier variant and falling within a 1Mb window around the QTL.

## Results

### High-quality reference assembly for *Coregonus clupeaformis* Normal species

Using long-read sequencing, we built the first reference genome assembly for *C. clupeaformis* sp. Normal (ASM1839867v1). The *de novo* assembly showed good contiguity with a N50 of 6.1 Mb and a L50 of 101 contigs. A linkage map allowed us to anchor and orient 83% of the genome into 40 linkage groups, the expected number of chromosomes for *C. clupeaformis* (Dion-Côté et al., 2015; Phillips, Reed, & Ráb, 1996), although some of the linkage groups, chromosome 22 in particular, may only represent a fraction of a chromosome. Studying recombination along those linkage groups, we identified 7 metacentric chromosomes, 3 putatively metacentric (or sub-metacentric chromosomes) and 30 acrocentric chromosomes (Fig. S4, Fig. 2A). The final assembly included 40 putative chromosomes and 6427 unanchored scaffolds with a N50 of 57 Mb for a total genome size of 2.68 Gb (Table 1). This reference genome had a high level of completeness, with 94% of universal single-copy orthologous genes in a BUSCO analysis based on the actonipterygii database. A relatively high percentage of duplicated BUSCO groups (44%) was observed, which is likely a consequence of the salmonid-specific whole genome duplication (Allendorf & Thorgaard, 1984; Smith et al., 2021).

**Figure 2:**
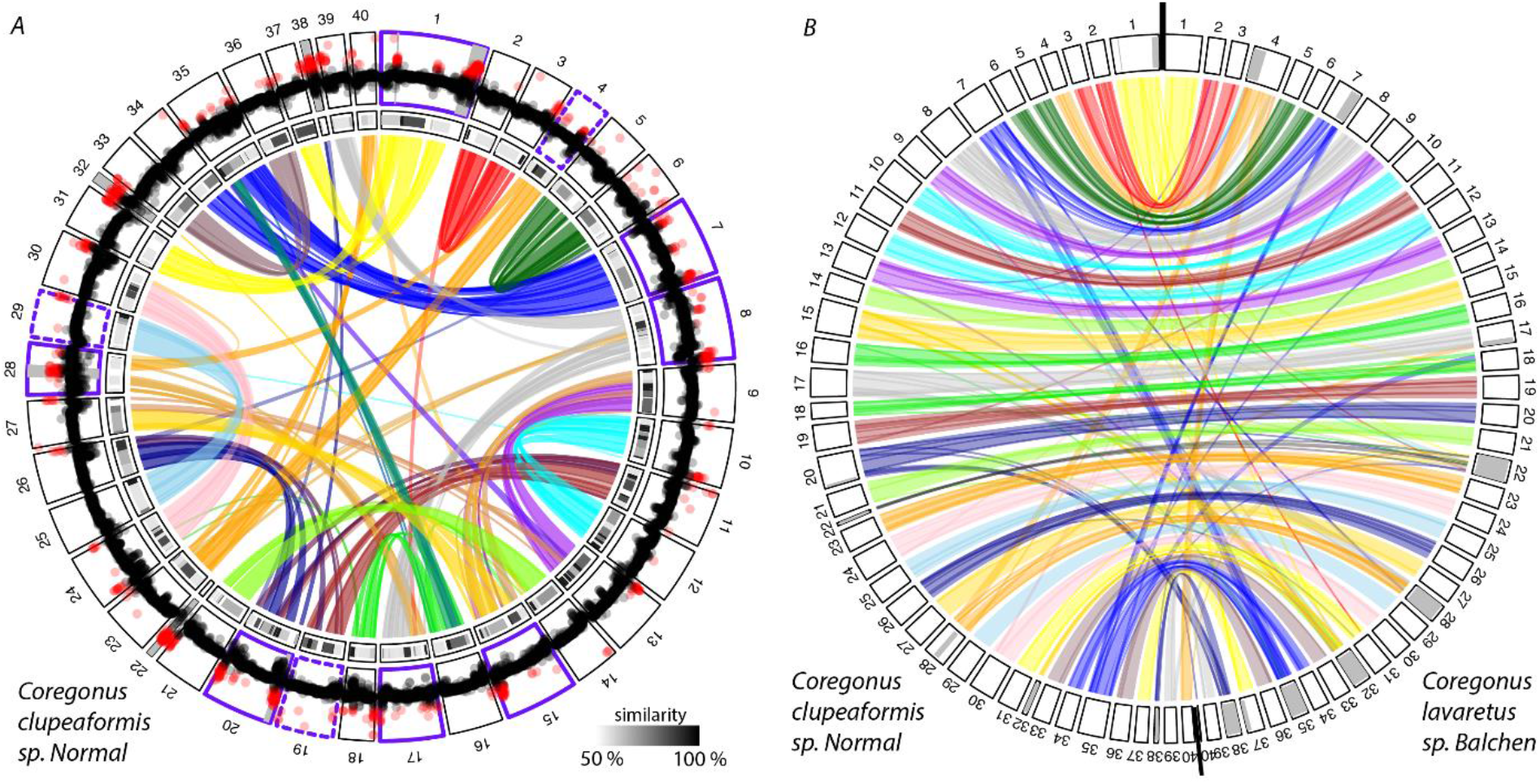
*Self-synteny and coverage in C. clupeaformis* sp. Normal *, and synteny with C. lavaretus* sp. *Balchen*. ***(A)*** Circular plot showing syntenic relationship between homoeologous chromosomes (inner track) and their level of sequence similarity (medium track) in the genome of *C. clupeaformis* sp. Normal. The outer track displays mean coverage by windows of 100kb in the short reads alignments. Points coloured in red show coverage higher than 1.5 times the average coverage (3.7X). Chromosomes surrounded by a purple outline are metacentric chromosomes, with dashed lines for putatively metacentric chromosomes. ***(B)*** Circular plot showing syntenic relationship between *C. clupeaformis* sp. Normal and *C. lavaretus* sp. *Balchen.* On both plots, chromosomes are coloured according to the ancestral origin (based on the PK nomenclature proposed in Sutherland et al (2016). Regions coloured in grey represent collapsed duplicated regions in genome assemblies.

**Table 1:** *Statistics of the genome assemblies of* C. clupeaformis *sp. Normal* and *sp. Dwarf.*

| Species | *C. clupeaformis* sp. Normal | *C. clupeaformis* sp. Dwarf |
| --- | --- | --- |
| Genome size | 2.68 Gb | 2.76 Gb |
| N50 (contig level) | 6.1 Mb | 2.2 Mb |
| L50 (contig level) | 101 contigs | 274 contigs |
| Fraction anchored in chromosomes | 83% | 73% |
| N50 (final assembly) | 57 Mb | 52 Mb |
| BUSCO score (Actinopterygii) | C:94.4% [S:50.9%, D:43.5%], F:1.7%, M:3.9%, n:4584 | C:94.6% [S:59.1%, D:35.5%], F:0.9%, M:4.5%, n:4584 |
| Fraction of TEs | 60.5% | 62.4% |

The genome of *C. clupeaformis* sp. Normal was composed of 60.5% of transposable elements (Fig. S5, Table S4). The greatest transposable element subclass representation in terms of total base pairs was DNA-TIR elements, taking up 24% of the genome. LINEs and LTRs were approximately equally abundant at about 13% of the genome each. Elements that were unclassified took up 9% of the genome. SINEs took up less than 1% of the genome, and rolling-circle/helitron elements were essentially absent. Our repeat identification pipeline identified 3490 distinct families. LTR elements were the most diverse with 1521 families identified, almost half the total number of families. Comparatively, 373 families were identified as DNA-TIR elements and 250 as LINEs. The genome of *C. clupeaformis* sp. Normal is composed of TEs at a variety of stages of decomposition (Fig. S6). A proxy for age of a given insertion is its sequence divergence from the consensus sequence, since the longer the insertions have been present, the more time there has been for accumulation of random mutations. The landscape plot shows that an equal amount (in terms of bp) of LINEs, LTRs, and DNA-TIRs are present in recent insertions less than 1% diverged from the consensus sequence. DNA-TIR elements near the 5% divergence level are the most abundant, indicating an older burst of activity.

The genome of *C. clupeaformis* sp. Normal showed high synteny with the closely-related European Alpine whitefish, *C. lavaretus “Balchen”* (Fig. 2A), allowing the identification of 39 homologous chromosomes which were named accordingly. Chromosome 40 of *C. lavaretus* sp. Balchen was small and had no homologous chromosome in genome of *C. clupeaformis* sp. Normal. Chromosome 40 of *C. clupeaformis* sp. Normal aligned with a fraction of chromosome 4 in the *C. lavaretus* sp. Balchen assembly and may or may not be one arm of the putatively-metacentric Chromosome 4. Some chromosomes (Chr7, Chr8, Chr15, Chr17, Chr20, Chr28, Chr35) included syntenic blocks matching two chromosomes in the related species. Some of those blocks likely correspond to duplicated regions collapsed in one of the assemblies, as they also exhibit higher than average coverage (Fig. 2A-2B). Those blocks may also belong to pseudo-tetrasomal chromosomes, which are homeologous chromosomes resulting from ancient whole-genome duplication and that still recombine to a certain extent (Allendorf et al., 2015; Blumstein et al., 2020; Lien et al., 2016; Waples, Seeb, & Seeb, 2016).

The identification of ancestral chromosomes by alignment to other linkage maps (Fig. S1-S3) and to the Northern Pike genome (Fig. S7), as well as self-synteny (Fig. 2A), allowed us to further identify the pairs of homeologous chromosomes. A few regions (Chr22, Chr 32, the end of Chr1) show no matching region in genome of *C. clupeaformis* sp. Normal but high coverage, suggesting that the assembly may have locally collapsed two highly similar regions (Fig. 2A, Table S3). Self-synteny assessment also supports fusion events between ancestral chromosomes that were previously reported in the three *Coregonus* species, *C. lavaretus, C. artedi* and *C. clupeaformis* (Blumstein et al., 2020; Sutherland et al., 2016) such as PK05-PK06 (Chr01), PK10-PK24 (chr8), PK11-PK21 (Chr7), PK01-PK14 (Chr15), PK16-PK23 (Chr4), PK8-PK9 (Chr20), as well as putative complex rearrangements between PK10-PK20-PK23 (Chr17, Chr28, Chr4). Those eight chromosomes also correspond to the ones identified as metacentric in our study and in the Cisco *C. artedii* (Blumstein et al., 2020).

### Discovery of SVs using a combination of sequencing tools

To identify SVs between Normal and Dwarf species, we built a *de novo* assembly for *C. clupeaformis* sp. Dwarf (ASM2061545v1) based on long-read sequences. This assembly shows high contiguity with a N50 of 2.2 Mb and a L50 of 274 contigs, of which 73% could be placed into chromosomes using the linkage map. The final Dwarf assembly included 40 chromosomes and 7294 unanchored scaffolds with a N50 of 52 Mb for a total genome size of 2.76 Gb. The Dwarf genome also showed high synteny with *C. lavaretus* sp. Balchen (Fig. S8). Like the Normal genome, the genome of *C. clupeaformis* sp. Dwarf was composed of about 61% of TE at various ages, with similar repartition between different class and families (Figure S5-S6, Table S6). The Dwarf genome also contains a high fraction (95%) of universal single-copy orthologous genes (actinopterygii), among which 36% were duplicated. This fraction is nevertheless smaller than in the Normal genome (44%), which may possibly reflect more collapsed duplicated regions in the Dwarf.

The comparison of the Dwarf assembly to the Normal reference unveiled a total of 244,717 SVs, of which 89,909 were detected by at least two tools and were kept as “high-confidence SVs”. Approximately half of the SVs were deletions and half were insertions (Fig. 3A). Duplications were counted as insertions, and only a limited number of inversions were detected (2,815, out of which only 77 were found by two tools).

**Figure 3:**
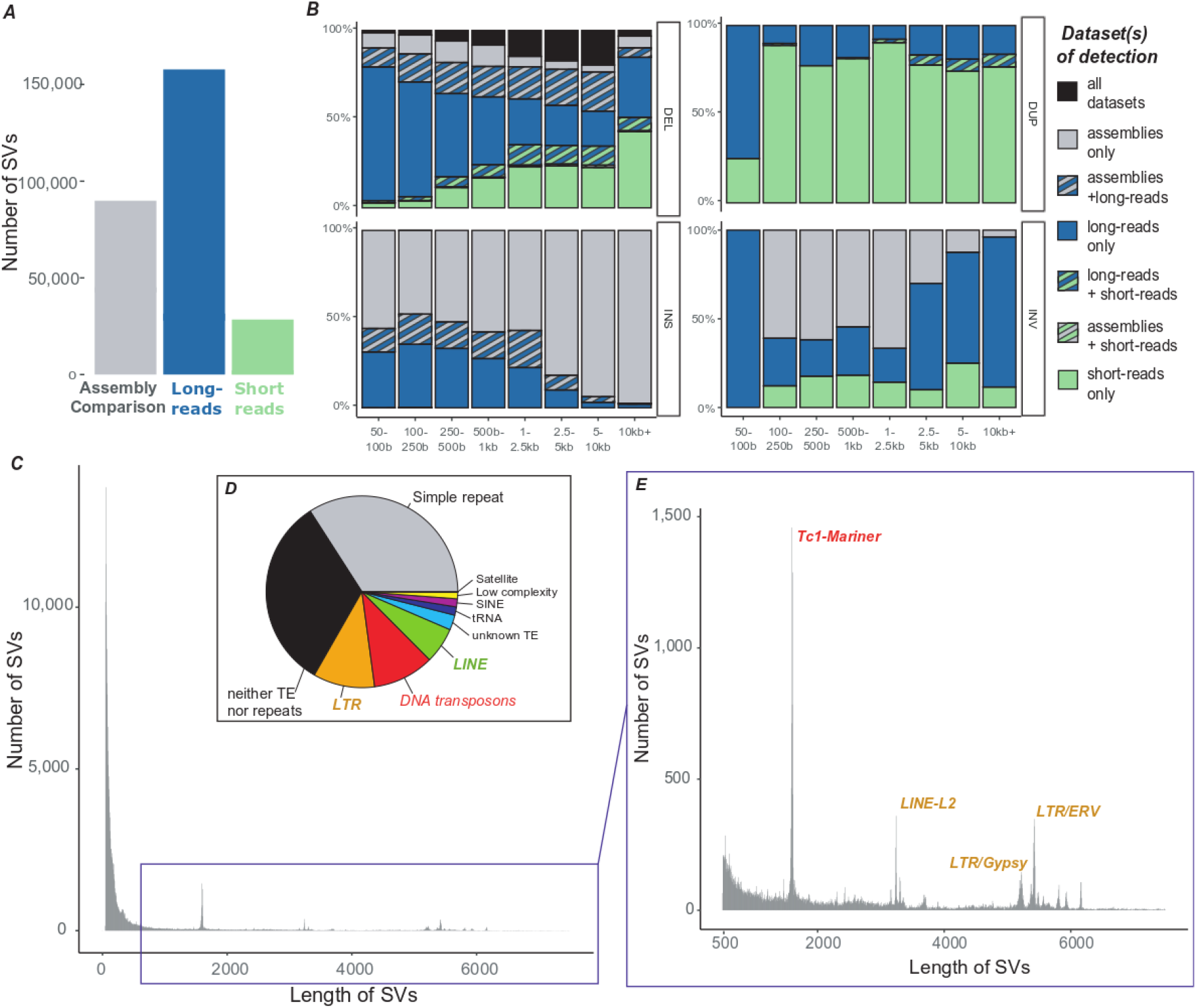
*Overview of SVs detected within and between C. clupeaformis* sp. Normal and sp. Dwarf. (A) Number of SVs detected in the three datasets by at least two tools (B) Proportion of SVs detected in one or several datasets according to type and size. (C) Size distribution of SVs. (D) Proportion of SVs associated with different families of transposable elements and repeated elements (E) Size distribution of SVs (zoomed on the range 500-7500bp).

Since a comparison of haploid assemblies is only able to detect SVs in the Dwarf relative to the Normal, and may be sensitive to assembly errors, we next investigated SV polymorphisms based on long reads. This unveiled a higher number of high-confidence SVs with a total of 194,861 SVs detected by at least two tools. Those included SVs putatively heterozygous in the Normal and the Dwarf genomes and resulted in a high number of novel deletions and insertions.

Only two samples (one Dwarf and one Normal) were sequenced with long reads; hence we hoped to cover a wider range of population structural polymorphism by using short reads on 32 individuals (15 Normal and 17 Dwarf) to detect SVs. This method nevertheless appeared less powerful than SV detection based on long-reads as 84,673 SVs were detected, with only 28,579 detected by at least two tools. This is possibly due to smaller size of short-reads and limited depth of sequencing in our dataset (about 5X), which is suboptimal for SV calling. The large majority of SVs detected in this dataset were deletions (n=77,899; 92%), followed by duplications (n=5,927;7%), a few inversions (n=24; 0.02%) and insertions (n=15; 0.01%) (Fig. 3A).

There was limited overlap between the different approaches with 7,525 SVs detected in the three datasets and 38,202 detected in two datasets out of a total of 222,927 SVs. This limited overlap, which varies depending on type and size, likely reflects the different sensitivity and detection power of the calling methods associated with each dataset. Almost no overlap was observed for inversions and duplications, likely reflecting the difficulties in characterizing such SVs. For insertions, the overlap between long reads and assembly comparison approaches tended to decrease with size, possibly due to more approximate breakpoints, while for deletions it increased with size (Fig. 3B).

The distribution of SV sizes was highly-skewed towards smaller SVs below 500bp (Fig. 3C). We observed heterogeneous peaks in the SV size distribution corresponding to insertions or deletions of transposable elements (Fig. 3E). The sequence of SVs around the 1600bp peak matches with TC1-Mariner. SVs around 3700bp correspond to Line-L2 indels while the peaks between 5000 and 6000bp are different kinds of LTR (Gypsy, ERV1). Overall, transposable elements were important factors driving SVs in *C. clupeaformis* as their sequences were composed of 73% of transposable elements (compared to 60% for the entire genome, Table S4). This enrichment was mostly due to retroelements (49% in SV sequences compared to 25 % in the genome), mostly LTR and Gypsy (Table S5). This resulted in about a third of all SVs in the catalog being associated with an insertion or deletion of a TE (Fig. 3D). Satellite repeats and simple repeats (e.g. microsatellites) cover a smaller fraction of the SV sequences (5%, Table S4) but they were found in about a third of SVs. A third of SVs did not match any TE nor any repeated regions.

### Polymorphism and differentiation in *C. clupeaformis* sp. Normal and sp. Dwarf

To assess genetic variation at the population level, we estimated genotype likelihoods for SNPs and SVs in the 32 samples sequenced with short reads. Filtering for genetic variants with allelic frequency higher than 5% retained 12,886,292 SNPs and 103,857 SVs. Those “frequent” SVs cover a total of 66 MB, representing polymorphism affecting approximately 5 times more nucleotides in the genome than SNPs. Decomposing genetic variation with a principal component analysis displayed a strong clustering of individuals by species and by lake. This was consistent whether considering SNPs or SVs, although SVs tended to show greater separation between the two species along the 1^st^ PC (Fig. 4A-B). This suggests a higher level of shared inter-specific variation between lakes for SVs than for SNPs.

**Figure 4:**
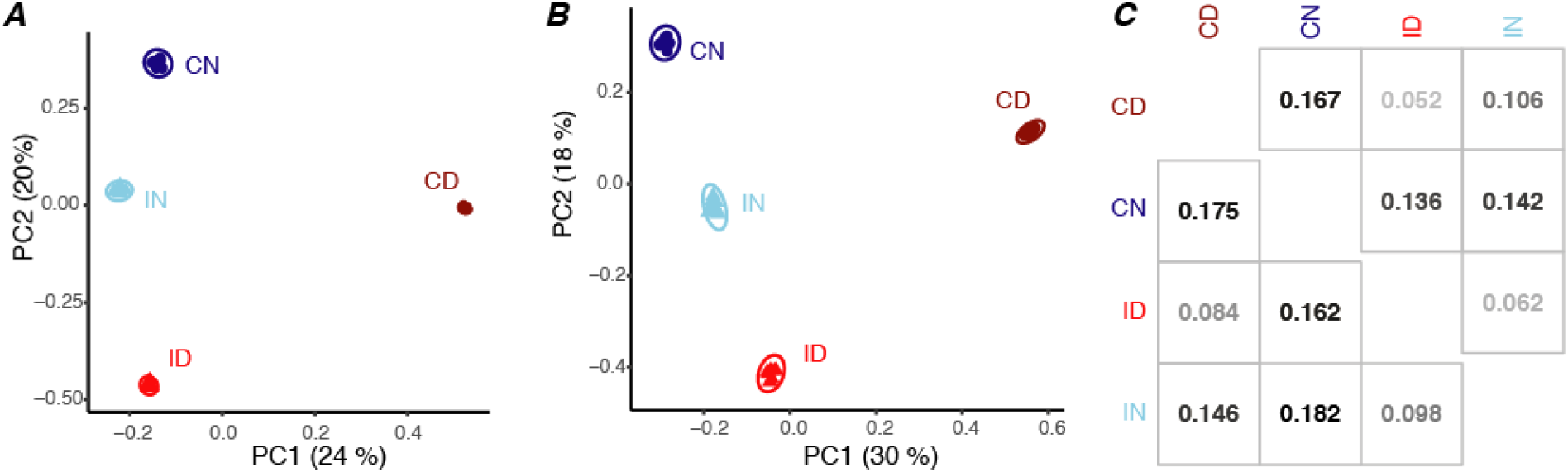
*Genomic variation in* C. clupeaformis *sp. Normal and sp. Dwarf*. Principal component analysis (PCA) based on (A) SNPs and (B) SVs. Each point is an individual coloured by lake and by species. (C) *F*_ST_ between lakes and species based on SNPs (below diagonal) and SVs (above diagonal). CN: Normal from Cliff Lake, CD: Dwarf from Cliff Lake, IN: Normal from Indian Lake, ID: Dwarf from Indian Lake

*F*_ST_ was moderate to high between lakes and between species, with values ranging from 0.052 up to 0.167 based on SVs and from 0.084 to 0.182 based on SNPs (Fig.4C). The Normal and Dwarf were more differentiated in Cliff Lake than in Indian Lake using both kinds of variants (Cliff Lake: *F*_ST_=0.175/0.167; Indian Lake: *F*_ST_=0.098/0.062) and such species differentiation was widespread along the genome (Fig. 5). Within each lake, the landscape of interspecific *F*_ST_ display similarities between SNPs and SVs, and 100kb window-based *F*_ST_ showed significant correlations when calculated on SNPs and on SVs (Cliff: R²=0.71, Indian: R²=0.63). This suggests that there may be linked variants (e.g., small deletions and SNPs) and that the two kinds of mutations may affect each other, for instance if some SVs reduce recombination.

**Figure 5:**
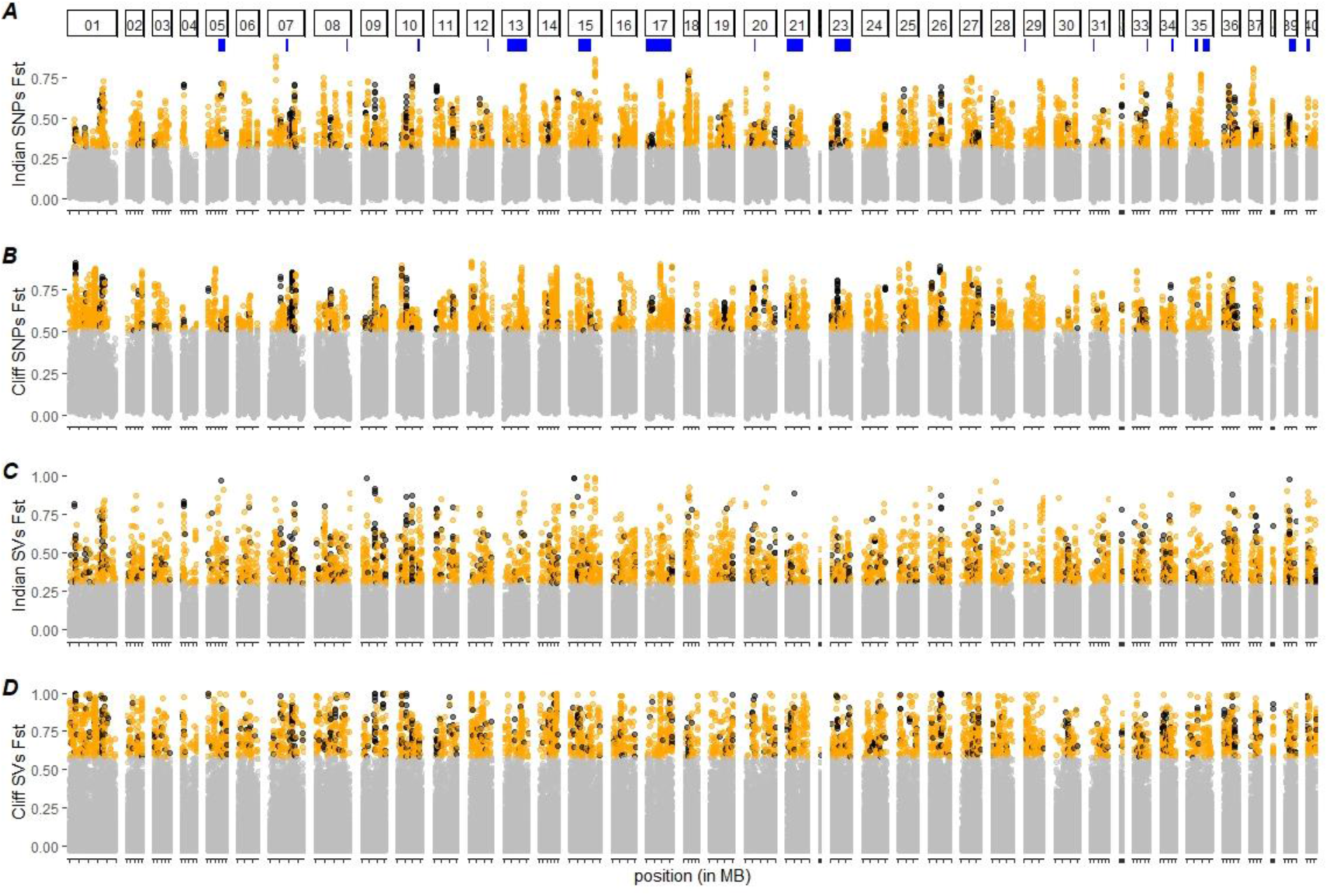
*Genomic differentiation along the genome between* C. clupeaformis *sp. Normal and* sp. *Dwarf*. *F*_ST_ between Normal and Dwarf based on SNPs, by windows of 100kb, in (A) Indian Lake and (B) Cliff Lake. *F*_ST_ between Normal and Dwarf based on SVs in (C) Indian Lake and (D) Cliff Lake. Windows and variants that exceed the 95% quantile in one lake are colored in orange. Shared polymorphisms between lakes (i.e., variants found as outliers in both lakes) are shown in black. Blue segments under chromosome numbers indicate the positions of QTLs associated with behavioral and morphological differences between Normal and Dwarf species, as identified in Gagnaire et al (2013).

As the two lakes represent parallel situations of coexistence between the Normal and the Dwarf species of *C. clupeaformis* (Rougeux et al., 2017), we investigated whether genetic differentiation follows similar patterns. The most differentiated genetic variants, defined as the SNPs and SVs in the top 95% *F*_ST_ quantile within each lake, showed three times the expected number of shared variants across lakes, suggesting that areas of differentiation between species are conserved in parallel across lakes. When measuring species differentiation as a polarized difference in allelic frequencies (AFD statistic), this overlap was even stronger. There was a fivefold excess for AFD outliers in the same end of the distribution (positive in both lakes and negative in both lakes). In other words, the variants with high allelic frequency differences between species are more likely than expected by chance to display the same Normal allele and Dwarf allele in both lakes (Table 2).

**Table 2:** Overlap across the two lakes in the most differentiated variants between species. *F*_ST_ outliers were defined as the top 5% of the *F*_ST_ distribution. AFD is the allelic frequency difference between the *C. clupeaformis* sp. Normal and *C. clupeaformis* sp. Dwarf (polarized as Dwarf-Normal). “Same sign” indicates that the outliers are in the same end of the AFD distribution in both lakes (either upper 97.5 quartile or lower 2.5 quartile), while “opposite sign” indicates that outliers are not in the same end of the AFD distribution in both lakes. In other words, outliers with opposite signs are variants in which the allele that is more frequent in the Dwarf in one lake is the allele that is more frequent in the Normal in the other lake.

| Dataset and method | Number of variants | Expected number of overlapping outliers | Observed number of overlapping outliers | Odds-ratio | p-value (Fisher test) |
| --- | --- | --- | --- | --- | --- |
| SNP $F_{ST}$ | 11,389,952 | 28,475 | 94,572 | 3.3 | $P<0.001$ |
| SNP AFD same sign | 11,389,952 | 14,237 | 80,474 | 5.7 | $P<0.001$ |
| SNP AFD opposite sign | 11,389,952 | 14,237 | 17,947 | 1.3 | $P<0.001$ |
| SV $F_{ST}$ | 93,773 | 234 | 727 | 3.1 | $P<0.001$ |
| SV AFD same sign | 93,773 | 117 | 618 | 5.3 | $P<0.001$ |
| SV AFD opposite sign | 93,773 | 117 | 123 | 1.1 | $P=0.75$ |

Relatively to all SVs, the most differentiated SVs, both within each lake and shared between lakes, were significantly enriched in TE-associated SVs. In other words, while SVs containing DNA transposons represent 15% of all SVs, they represent 27% of outlier SVs. In contrast, SVs associated with simple repeats were underrepresented in outliers of differentiation, while SVs without TE or repeats showed no bias. This excess of TE-linked SVs in outliers was driven by all categories of TE: DNA transposons, LINEs, SINEs and LTR. The most significantly enriched families in both lakes were the DNA transposons Tc1-mariner and hAT-Ac, and the retrotransposons LTR-Gypsy and LTR-ERV1, LINE-L1, LINE-L2 and LINE-RexBabar (Table 3, Table S7).

**Table 3:**
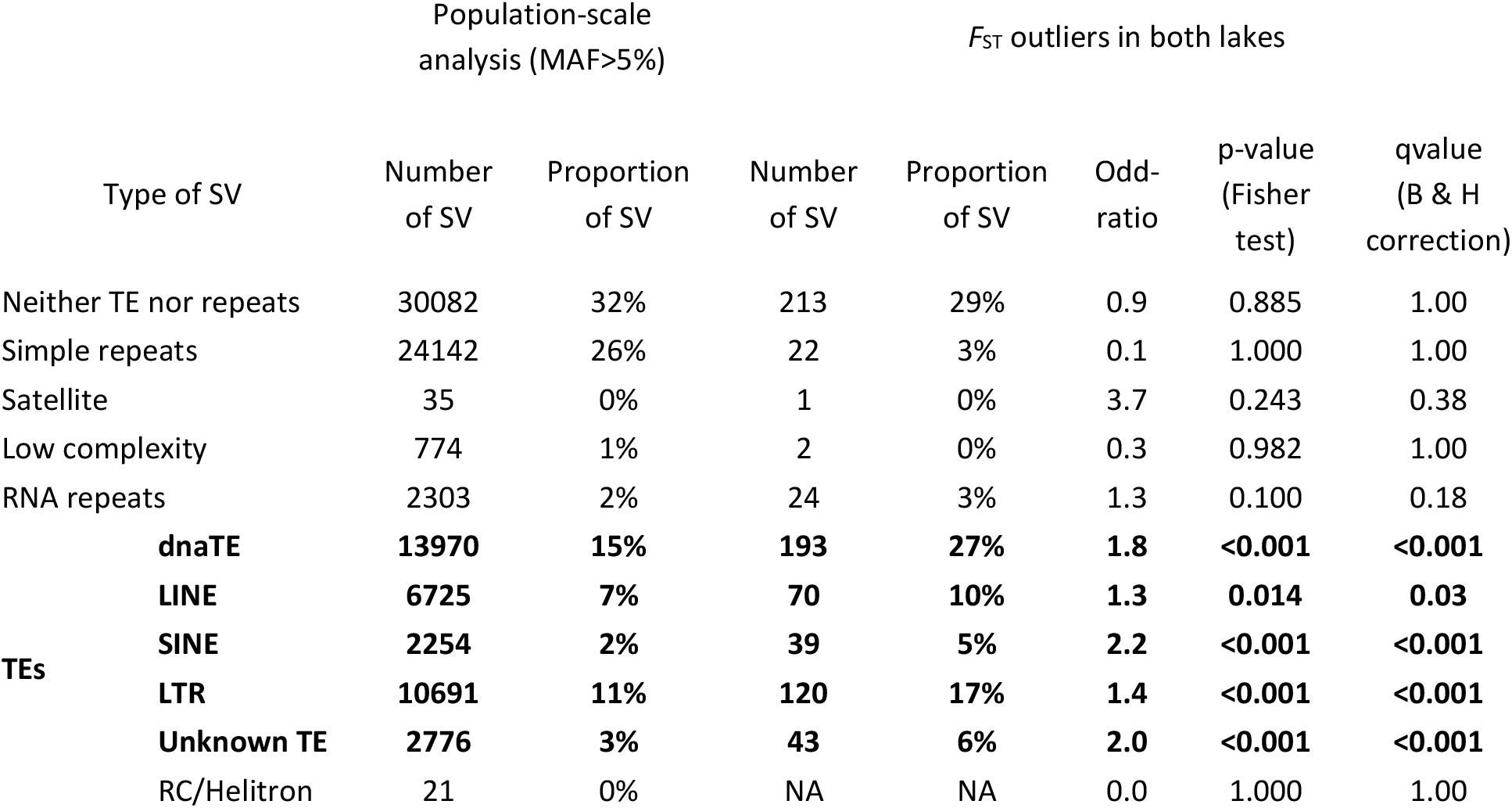
*Enrichment in SVs associated with transposable elements in outliers of differentiation between* C. clupeaformis *sp. Normal and* C. clupeaformis *sp. Dwarf* .

The most differentiated variants overlapped with thousands of genes. Out of a total of 34,913 genes with SNPs, 15,732 genes had at least one outlier SNP in Cliff Lake, 17,344 in Indian Lake, and 4,678 in both lakes. Out of a total of 13,886 genes with SVs, 1,396 genes had at least one outlier SV in Cliff Lake, 1,622 in Indian Lake, and 242 in both lakes. Gene ontology analysis revealed significant enrichment in behavior, morphogenesis, cell signaling, immunity and metabolism, among many other fucntions (Table S8). To narrow down putative candidate genes possibly involved in phenotypic differentiation, we focused on outliers overlapping with QTLs previously mapped with the linkage map (Gagnaire, Normandeau, et al., 2013; S. M. Rogers et al., 2007). A total of 27 QTLs for various traits differentiating Dwarf and Normal (growth rate, maturity, gill raker, etc) could be positioned on the new reference genome, although some of them had a relatively wide interval (Table S9, Fig 5). They overlapped with 45,823 SNPs and 414 SVs that were identified as outliers of differentiation in both lakes. The list of genes belonging to a QTL and overlapping with at least one outlier is provided in Table S10.

## Discussion

By combining long-read and short-read sequencing on two species of the Lake Whitefish complex, *C. clupeaformis* sp. Normal and *C. clupeaformis* sp. Dwarf, our study generated new genomic resources and provided insights into the genomic architecture of recent speciation. First, we have produced a reference genome assembly for both *C. clupeaformis* sp. Normal and *C. clupeaformis* sp. Dwarf, as well as an extensive catalog of SVs. Second, studying SVs at the population-level showed that SVs represent a large amount of variation within and between Normal and Dwarf sympatric species, less numerous but more extensive than SNPs in terms of total number of nucleotides. Third, by comparing young species pairs in two lakes, we highlighted shared genetic differentiation and supported a predominant role of transposable elements in the divergence between the Normal and the Dwarf. Hereafter, we discuss how our results and methods contribute to a better understanding of the genomic architecture of speciation and the involvement of structural polymorphism.

Generating high-quality reference genomes for non-model species is becoming a requirement to understand the evolution of genomic variation during the speciation process (Nadeau & Kawakami, 2019; Ravinet et al., 2017). Here, using the genome of *C. clupeaformis* sp. Normal from North America as reference facilitated the accurate detection of population-level variants, both SNPs and SVs, because the reference is from the same species, or a closely-related species, and from the same geographic area. Moreover, contiguity and chromosome-level information allowed a finer understanding of the role played by recombination, large rearrangements and chromosome-level variability (fusion/fission, karyotypic polymorphism, etc). In our study, the use of long-reads proved incomparable to resolve the complexity of the genomes of *C. clupeaformis* sp. Normal and sp. Dwarf. Using Nanopore sequencing data, we have been able to reach a high contiguity, allowing us to search for SVs. Assembly comparison, as well as direct detection based on long-reads, show that one Normal and one Dwarf individuals differ by more than 100,000 high-confidence SVs. Given the stringency of our quality filters, and the lack of power to detect complex rearrangements or inversions, this number should be seen as a lower bound of the amount of SVs. In particular, most of the detected SVs remain in a range of small size (<1kb) or relatively medium size. This catalog of SVs can therefore be supplemented by including more individuals, longer sequences, and additional genomes. In any case, the large number of high-confidence SVs identified in this study reinforces the importance of considering the possible role of SVs in evolutionary processes such as adaptation and speciation.

Regarding SVs of larger size (>100kb), we acknowledge that the detection power of our dataset was limited. Because the final scaffolding of the two genomes is based on a single (and not so dense) linkage map, made from a Normal × Dwarf hybrid family (Gagnaire, Normandeau, et al., 2013; S. M. Rogers et al., 2007), we could not search for large chromosomal rearrangements simply by contrasting the two genomes. This is unfortunate since large rearrangements such as inversions, fusions, and translocations may be relevant for speciation because they often differ between closely-related sympatric species and contribute to reproductive isolation (Berdan et al., 2021; Faria & Navarro, 2010; Noor et al., 2001). In the case of *C. clupeaformis*, on the one hand, we do not expect a major effect of chromosomal rearrangements. First, the differentiation observed in SNPs and SVs is widespread along the genome and does not display the typical spatial clustering of differentiated regions observed between species pairs like *Littorina saxatilis* (Morales et al., 2019) or *Helianthus* sp (Todesco et al., 2020). Second, cytogenetic analysis showed that the *C. clupeaformis* sp. Normal and sp. Dwarf from these same lakes have an identical number of chromosomes (Dion-Côté et al., 2017). On the other hand, cytogenetic exploration showed subtle chromosomal polymorphism within and between them (Dion-Côté et al., 2017). For instance, chromosome 1 is longer in the Normal than in the Dwarf in Cliff Lake due to heterochromatin differences (Dion-Côté et al., 2017), a pattern that we also observed in the genome (121Mb vs. 99Mb, Fig. S8). We also note some peculiarities such as Chr22, for which sequences in *C. clupeaformis* sp. Normal are homologous to sequences belonging to chromosome 22 in the genome of *C. lavaretus* sp. Balchen but which we never managed to order as a full linkage group, likely because of the lack of recombination in the family used for the linkage map. Since the mother used for the linkage map is a hybrid Dwarf × Normal, any rearrangement differing between species (and affecting recombination at heterozygote stage) may be absent from the final map, and hence from the present genomes. These chromosomal differences may lead to issues with recombination during meiosis (Dion-Côté et al., 2015; Faria & Navarro, 2010), contributing to reproductive isolation and speciation (Hoffmann & Rieseberg, 2008; Kirkpatrick & Barton, 2006). In the future, it would be worthwhile to explore large scale chromosomal rearrangements in *C. clupeaformis* sp. Normal and *C. clupeaformis* sp. Dwarf in depth to understand the role of chromosomal polymorphism in speciation. However, this will require improved genome scaffolding based on Hi-C chromatin contacts (which was attempted here without success) or separate linkage maps.

Beyond the contrast between *C. clupeaformis* sp. Normal and *C. clupeaformis* sp. Dwarf, the new genome assemblies also provide relevant information about the evolution of genomes at a higher taxonomic level. Salmonids have experienced a recent whole-genome duplication, followed by different events of re-diploidization, as well as important chromosomal rearrangements such as fusions (Blumstein et al., 2020; Glasauer & Neuhauss, 2014; Lien et al., 2016; Macqueen & Johnston, 2014). Here, as often observed in salmonids, synteny was high between *C. clupeaformis* sp. Normal, *C. clupeaformis* sp. Dwarf, and closely related species such as the European whitefish *C. lavaretus* sp. Balchen. The same groups of chromosomes appear to be metacentric and bear residual tetrasomy in *C. clupeaformis* as in its related species *C. ardetii* (Blumstein et al., 2020). Chromosomal comparison with *C. ardetii* and *C. lavaretus* also suggested shared fusion and fission of ancestral chromosomes and a consistent karyotype between the different coregonids (Blumstein et al., 2020; De-Kayne & Feulner, 2018). This would suggest that the majority of re-deploidization processes occurred before the split of the different *Coregonus* species, which would all share a relatively similar karyotype. That being said, it should be kept in mind that the residual tetrasomy observed on a subset of chromosomes makes it difficult to fully ascertain synteny vs. rearrangements within and between species on those chromosomes. Moreover, *C. clupeaformis* genomes remain extremely complex with several regions that end up collapsed by genome assembly (at least 126 Mb, 5% of the chromosome-anchored genome), as was previously reported in other salmonid genomes (De-Kayne et al., 2020; Lien et al., 2016). Therefore, while the *Coregonus* reference genome assemblies provide an important first step, refining the assemblies and complementing by cytogenetic or chromatin-contact data will be valuable to further explore of the timing and modalities of re-diploidization in coregonids, and its possible contribution to speciation.

Salmonids genomes are also littered with transposable elements and *C. clupeaformis* was no exception: interspersed repeats accounted for about 60% of the genome. This amount is comparable to *Salmo salar* (60% (Lien et al. 2016)) and *Coregonus lavaretus* “Balchen” (52% (De-Kayne et al. 2020)). Moreover, all TE copies are not shared by all individuals and our results highlighted that they were responsible for a third of the SVs detected within and between species. This is also consistent with observations made on other species, such as Atlantic Salmon *Salmo salar* (Bertolotti et al., 2020) or songbird *Corvus* sp. (Weissensteiner et al., 2020), in which young and active TEs generate numerous insertions and deletions between samples. It has been hypothesized that bursts of transposon activity may contribute to speciation (de Boer, Yazawa, Davidson, & Koop, 2007), or at least that TEs may rapidly generate genetic variation differentiating species (Serrato-Capuchina & Matute, 2018). Our data strongly support this hypothesis since the most differentiated SVs between Dwarf and Normal in both lakes were enriched in several classes of TEs. A large part of the fixed genetic variation between species corresponds to an insertion or a deletion of a given TE. It is worth noting that this pattern is widespread across the genome rather than centered on a few loci. Such extensive differentiation suggests a progressive and differential TE accumulation without gene flow, likely in allopatry during the Pleistocene glaciation (~15,000 generations/60,000 years ago) that may have contributed to the maintenance of reproductive isolation during the postglacial sympatric phase following secondary contact (~3,000 generations/12,000 years ago) (Jacobsen et al., 2012; Rougeux et al., 2017). Accumulations of different TEs between lineages may be quite rapid as active TEs have a high mutation rate, as observed in Daphnia with an order of 10^−5^ gain or loss per copy per generation (Ho et al., 2021). TEs can also contribute to reproductive isolation by altering gene structure, expression pattern and chromosome organization (Dubin, Scheid, & Becker, 2018; Goodier, 2016). In fact, TE deregulation is known to generate post-zygotic breakdown in Dwarf × Normal hybrids (Dion-Côté, Renaut, Normandeau, & Bernatchez, 2014), which has been associated with epigenetic (DNA methylation) reprogramming in hybrids (Laporte et al., 2019). Moreover, this supported the hypothesis that TE transcriptional derepression, perhaps due to different TE silencing mechanisms that evolved in allopatry, may be the cause for both massive misregulation of gene expression and abnormal embryonic development and death in hybrids (Renaut, Nolte, & Bernatchez, 2009; Dion-Côté et al., 2014). Both in previous studies and in our study, the same TE families emerged as associated with species differentiation, namely Tc1-mariner and hAT-Ac as well as LTR-Gypsy, Line-L2 and Line-RexBabar. Altogether, cumulative evidence points towards a major role of several TE families in the reproductive isolation of Dwarf and Normal, involving TEs distributed throughout the genome rather than in a few barrier loci.

A peculiarity of the speciation between *C. clupeaformis* sp. Normal and *C. clupeaformis* sp. Dwarf is the character displacement in the Acadian lineage towards a dwarf limnetic species upon secondary contact with the American lineage, a process with occurred independently in separate lakes of the suture zone, resulting in two ecologically distinct sympatric species, the Dwarf and the Normal (Bernatchez et al., 2010a; Landry et al., 2007; Rougeux et al., 2017) . Previous work revealed that strong parallelism at the phenotypic level between lakes was accompanied by weak parallelism at the genome level (Gagnaire, Pavey, et al., 2013; Lu & Bernatchez, 1999; Rougeux, Gagnaire, Praebel, Seehausen, & Bernatchez, 2019). With a higher density of variants being screened, our results corroborate those from these previous studies. The pattern of differentiation between species was indeed specific to each lake. However, it is worth noting the excess of shared outliers of differentiation, both for SNPs and SVs, and that differences of allelic frequencies were more often in the same direction (e.g., higher allelic frequency in dwarf species in both lakes) than expected by chance. A large fraction of such parallelism likely reflects historical divergence between allopatric lineages, possibly reinforced by the result of comparable ecological response to selection. It is also possible that shared regions of differentiation reflect regions of the genome more resistant to gene flow, such as low recombination regions, as observed in *Ficedula* flycatchers (Burri et al., 2015). General patterns of TE enrichment in outlier SVs, as well as gene ontology enrichment, also converged between lakes. This suggests that the processes driving genetic divergence between species were likely similar between lakes, namely through shared historical divergence and similar ecological selection imposed by the use of distinct trophic niches (Bernatchez et al., 2010a). However, they were buffered by lake-specific contingency at finer molecular level, for instance, associated with the effect of genetic drift on available standing genetic variation within each lake (Gagnaire, Pavey, et al., 2013).

Studying two types of genetic variants in tandem, SVs and SNPs, at the population level showed similar patterns and level of differentiation between species and between lakes. On the one hand, this confirms that evaluating population/species structure requires neither a diversity of variants nor a large amount of markers. In fact, the *F*_ST_ values observed at the scale of the entire genome for both types of variants and in both lakes were strikingly similar to values measured based on a much smaller subset of markers. For instance, based on the RADseq genotyping of about 2500 SNP loci, Gagnaire, Pavey, et al. (2013) reported *F*_ST_ values of 0.12 and 0.10 between Dwarf and Normal from Cliff Lake and from Indian Lake respectively, compared to values of 0.18 and 0.10 here for SNPs and 0.17 and 0.06 for SVs. On the other hand, studying different kinds of variants with similar filters shows a large amount of non-rare SVs, *i.e.* SVs found in more than 2 of 64 chromosomes (32 diploid individuals). Because of their size, the accumulation of SVs at intermediate frequency in natural populations thus represents a non-negligible aspect of genetic variation, as they covered at least five times more of the genome than SNPs. This point is increasingly underlined by studies in population genomics and evolutionary genomics (Catanach et al., 2019; Mérot et al., 2020; Weissensteiner et al., 2020) and means that a full understanding of genetic variation cannot overlook SVs. However, it remains difficult to study SVs at the population level. Short reads are more accessible when sequencing a large number of individuals but they proved to be less powerful for characterizing SVs (Mahmoud et al., 2019). For instance, here we found around 5 times fewer SVs with 32 samples sequenced with short-reads than with 2 samples sequenced with long reads. Our study also used short-reads at shallow/medium coverage (~5X) which may be suboptimal to detect and genotype SNPs and SVs with confidence. However, there are ways to handle the uncertainty associated with a low number of supporting reads, such as working within a genotype likelihood framework (Alex Buerkle & Gompert, 2013; Lou et al., 2020). Recent studies have proposed relying on mixed datasets (e.g. combining long-read and short-read sequence data, combining high and shallow coverage) to achieve together a good catalog of SVs and then perform population genomic studies based on their variation (Logsdon et al., 2020). We have achieved this in this study by first characterizing SVs using high-depth long reads on a limited number of samples, and secondly by genotyping known SVs with medium-coverage short-reads on a greater number of samples. To achieve this, genome-graph based approaches were particularly relevant, allowing us to build a variation-aware reference graph (Garrison et al., 2018), and then perform unbiased mapping of reads to this graph (Sirén et al., 2020). Such two-step approaches have also be used in a handful of studies looking at SVs in chocolate trees *Theobroma cacao* (Hämälä et al., 2021), soybeans *Glycine max* (Lemay et al., 2021), and potato beetle *Leptinotarsa decemlineata* (Cohen, Hawthorne, & Schoville, 2021). Based on this, we believe that the combination of 2^nd^ and 3^rd^ generation sequencing is promising to study structural polymorphism within a population genomics framework and will allow the inclusion of SVs in studies of speciation and adaptation genomics.

## Supporting information

TabS8

TabS10

## Acknowledgements

We would like to thank M. Suga for contributing to fieldwork, K. Wellband for help with annotation, M-A. Lemay for help with SV analysis, M. Leitwein for help with the analysis of metacentric chromosomes, and A-L Ferchaud and R. De Kayne for their help with synteny analysis and plotting. We are very grateful to the teams that developed vg, giraffe and pggb for their guidance in using the genome-graph tools as well as J. Laroche at the IBIS Bioinformatic platform (www.ibis.ulaval.ca) for his support. We are also grateful to Editor L. Rieseberg and three anonymous reviewers for their extensive and constructive comments on a previous version of this manuscript.

## Data Availability

Final genome assemblies are available on NCBI under the id ASM1839867v1 and ASM2061545v1. Long-read sequences and short-read sequences will be available on NCBI under project number XXX upon publication. Code are available on Github for the different pipelines listed in the methods.

## Author contributions

This study is part of the long-term research program of L.B. on the adaptive radiation and ecological speciation of Lake Whitefish. L.B., C.R. and C.M. conceived the study. M.Á. and M.K. performed the Nanopore sequencing. M.M and S.L. assembled the genome and E.N. and C.M. scaffolded the genome. J.M.F. analysed the transposable elements. C.M. and K.S. did the analyses with contributions from C.V., E.N., M.L. and S.L. C.M. wrote the manuscript with contributions from all authors.

## References

Abel, H. J., Larson, D. E., Regier, A. A., Chiang, C., Das, I., Kanchi, K. L., … Reeves, C. (2020). Mapping and characterization of structural variation in 17,795 human genomes. Nature, 583(7814), 83–89.

Alex Buerkle, C., & Gompert, Z. (2013). Population genomics based on low coverage sequencing: How low should we go? Molecular Ecology, 22(11), 3028–3035.

Aljanabi, S. M., & Martinez, I. (1997). Universal and rapid salt-extraction of high quality genomic DNA for PCR-based techniques. Nucleic Acids Research, 25(22), 4692–4693.

Allendorf, F. W., Bassham, S., Cresko, W. A., Limborg, M. T., Seeb, L. W., & Seeb, J. E. (2015). Effects of crossovers between homeologs on inheritance and population genomics in polyploid-derived salmonid fishes. Journal of Heredity, 106(3), 217–227.

Allendorf, F. W., & Thorgaard, G. H. (1984). Tetraploidy and the evolution of salmonid fishes. In Evolutionary genetics of fishes (pp. 1–53). Springer.

Alonge, M., Soyk, S., Ramakrishnan, S., Wang, X., Goodwin, S., Sedlazeck, F. J., … Schatz, M. C. (2019). RaGOO: fast and accurate reference-guided scaffolding of draft genomes. Genome Biology, 20(1), 1–17.

Berdan, E. L., Fuller, R. C., & Kozak, G. M. (2021). Genomic landscape of reproductive isolation in Lucania killifish: The role of sex loci and salinity. Journal of Evolutionary Biology, 34(1), 157–174.

Bernatchez, L., & Dodson, J. J. (1990). Allopatric origin of sympatric populations of lake whitefish (Coregonus clupeaformis) as revealed by mitochondrial-DNA restriction analysis. Evolution, 44(5), 1263–1271.

Bernatchez, L., Renaut, S., Whiteley, A. R., Derome, N., Jeukens, J., Landry, L., … Rogers, S. M. (2010a). On the origin of species: Insights from the ecological genomics of lake whitefish. Philosophical Transactions of the Royal Society B: Biological Sciences, 365(1547), 1783–1800.

Bernatchez, L., Renaut, S., Whiteley, A. R., Derome, N., Jeukens, J., Landry, L., … Rogers, S. M. (2010b). On the origin of species: Insights from the ecological genomics of lake whitefish. Philosophical Transactions of the Royal Society B: Biological Sciences, 365(1547), 1783–1800.

Bertolotti, A. C., Layer, R. M., Gundappa, M. K., Gallagher, M. D., Pehlivanoglu, E., Nome, T., … Macqueen, D. J. (2020). The structural variation landscape in 492 Atlantic salmon genomes. Nature Communications, 11(1), 5176. doi: 10.1038/s41467-020-18972-x

Blumstein, D. M., Campbell, M. A., Hale, M. C., Sutherland, B. J., McKinney, G. J., Stott, W., & Larson, W. A. (2020). Comparative genomic analyses and a novel linkage map for cisco (Coregonus artedi) provide insights into chromosomal evolution and rediploidization across salmonids. G3: Genes, Genomes, Genetics, 10(8), 2863–2878.

Breese, M. R., & Liu, Y. (2013). NGSUtils: A software suite for analyzing and manipulating next-generation sequencing datasets. Bioinformatics, 29(4), 494–496.

Burri, R., Nater, A., Kawakami, T., Mugal, C. F., Olason, P. I., Smeds, L., … Garamszegi, L. Z. (2015). Linked selection and recombination rate variation drive the evolution of the genomic landscape of differentiation across the speciation continuum of Ficedula flycatchers. Genome Research, 25(11), 1656–1665.

Cabanettes, F., & Klopp, C. (2018). D-GENIES: dot plot large genomes in an interactive, efficient and simple way. PeerJ, 6, e4958.

Catanach, A., Crowhurst, R., Deng, C., David, C., Bernatchez, L., & Wellenreuther, M. (2019). The genomic pool of standing structural variation outnumbers single nucleotide polymorphism by threefold in the marine teleost Chrysophrys auratus. Molecular Ecology, 28(6), 1210–1223.

Catchen, J., Amores, A., & Bassham, S. (2020). Chromonomer: A tool set for repairing and enhancing assembled genomes through integration of genetic maps and conserved synteny. BioRxiv.

Chaisson, M. J., Sanders, A. D., Zhao, X., Malhotra, A., Porubsky, D., Rausch, T., … Collins, R. L. (2019). Multi-platform discovery of haplotype-resolved structural variation in human genomes. Nature Communications, 10(1), 1–16.

Chakraborty, M., Baldwin-Brown, J. G., Long, A. D., & Emerson, J. (2016). Contiguous and accurate de novo assembly of metazoan genomes with modest long read coverage. Nucleic Acids Research, 44(19), e147–e147.

Chen, S., Zhou, Y., Chen, Y., & Gu, J. (2018). fastp: An ultra-fast all-in-one FASTQ preprocessor. Bioinformatics, 34(17), i884–i890.

Chen, X., Schulz-Trieglaff, O., Shaw, R., Barnes, B., Schlesinger, F., Källberg, M., … Saunders, C. T. (2016). Manta: Rapid detection of structural variants and indels for germline and cancer sequencing applications. Bioinformatics, 32(8), 1220–1222.

Cohen, Z., Hawthorne, D., & Schoville, S. (2021). The role of structural variants in pest adaptation and genome evolution of the Colorado potato beetle, Leptinotarsa decemlineata (Say). Authorea Preprints.

Cruickshank, T. E., & Hahn, M. W. (2014). Reanalysis suggests that genomic islands of speciation are due to reduced diversity, not reduced gene flow. Molecular Ecology, 23(13), 3133–3157.

Dalziel, A. C., Laporte, M., Rougeux, C., Guderley, H., & Bernatchez, L. (2017). Convergence in organ size but not energy metabolism enzyme activities among wild Lake Whitefish (Coregonus clupeaformis) species pairs. Molecular Ecology, 26(1), 225–244.

de Boer, J. G., Yazawa, R., Davidson, W. S., & Koop, B. F. (2007). Bursts and horizontal evolution of DNA transposons in the speciation of pseudotetraploid salmonids. BMC Genomics, 8(1), 1–10.

De Coster, W., De Rijk, P., De Roeck, A., De Pooter, T., D’Hert, S., Strazisar, M., … Van Broeckhoven, C. (2019). Structural variants identified by Oxford Nanopore PromethION sequencing of the human genome. Genome Research, 29(7), 1178–1187.

De-Kayne, R., & Feulner, P. G. (2018). A European whitefish linkage map and its implications for understanding genome-wide synteny between salmonids following whole genome duplication. G3: Genes, Genomes, Genetics, 8(12), 3745–3755.

De-Kayne, R., Zoller, S., & Feulner, P. G. (2020). A de novo chromosome-level genome assembly of Coregonus sp.“Balchen”: One representative of the Swiss Alpine whitefish radiation. Molecular Ecology Resources, 20(4), 1093–1109.

Dion-Côté, A., Symonová, R., Lamaze, F. C., Pelikánová, Š., Ráb, P., & Bernatchez, L. (2017). Standing chromosomal variation in Lake Whitefish species pairs: The role of historical contingency and relevance for speciation. Molecular Ecology, 26(1), 178–192.

Dion-Côté, A.-M., Renaut, S., Normandeau, E., & Bernatchez, L. (2014). RNA-seq reveals transcriptomic shock involving transposable elements reactivation in hybrids of young lake whitefish species. Molecular Biology and Evolution, 31(5), 1188–1199.

Dion-Côté, A.-M., Symonová, R., Ráb, P., & Bernatchez, L. (2015). Reproductive isolation in a nascent species pair is associated with aneuploidy in hybrid offspring. Proceedings of the Royal Society B: Biological Sciences, 282(1802), 20142862.

Dubin, M. J., Scheid, O. M., & Becker, C. (2018). Transposons: A blessing curse. Current Opinion in Plant Biology, 42, 23–29.

Dufresnes, C., Brelsford, A., Jeffries, D. L., Mazepa, G., Suchan, T., Canestrelli, D., … Crochet, P.-A. (2021). Mass of genes rather than master genes underlie the genomic architecture of amphibian speciation. Proceedings of the National Academy of Sciences, 118(36), e2103963118. doi: 10.1073/pnas.2103963118

Evans, M., & Bernatchez, L. (2012). Oxidative phosphorylation gene transcription in whitefish species pairs reveals patterns of parallel and nonparallel physiological divergence. Journal of Evolutionary Biology, 25(9), 1823–1834.

Faria, R., & Navarro, A. (2010). Chromosomal speciation revisited: Rearranging theory with pieces of evidence. Trends in Ecology & Evolution, 25(11), 660–669.

Feulner, P., Chain, F. J., Panchal, M., Eizaguirre, C., Kalbe, M., Lenz, T. L., … Milinski, M. (2013). Genome-wide patterns of standing genetic variation in a marine population of three-spined sticklebacks. Molecular Ecology, 22(3), 635–649.

Feulner, P., & De-Kayne, R. (2017). Genome evolution, structural rearrangements and speciation. Journal of Evolutionary Biology, 30(8), 1488–1490.

Flynn, J. M., Hubley, R., Goubert, C., Rosen, J., Clark, A. G., Feschotte, C., & Smit, A. F. (2020). RepeatModeler2 for automated genomic discovery of transposable element families. Proceedings of the National Academy of Sciences, 117(17), 9451–9457.

Gagnaire, P., Normandeau, E., Pavey, S. A., & Bernatchez, L. (2013). Mapping phenotypic, expression and transmission ratio distortion QTL using RAD markers in the Lake Whitefish (Coregonus clupeaformis). Molecular Ecology, 22(11), 3036–3048.

Gagnaire, P., Pavey, S. A., Normandeau, E., & Bernatchez, L. (2013). The genetic architecture of reproductive isolation during speciation-with-gene-flow in lake whitefish species pairs assessed by RAD sequencing. Evolution, 67(9), 2483–2497.

Garrison, E., Sirén, J., Novak, A. M., Hickey, G., Eizenga, J. M., Dawson, E. T., … Lin, M. F. (2018). Variation graph toolkit improves read mapping by representing genetic variation in the reference. Nature Biotechnology, 36(9), 875–879.

Glasauer, S. M., & Neuhauss, S. C. (2014). Whole-genome duplication in teleost fishes and its evolutionary consequences. Molecular Genetics and Genomics, 289(6), 1045–1060.

Goel, M., Sun, H., Jiao, W.-B., & Schneeberger, K. (2019). SyRI: finding genomic rearrangements and local sequence differences from whole-genome assemblies. Genome Biology, 20(1), 1–13.

Goodier, J. L. (2016). Restricting retrotransposons: A review. Mobile DNA, 7(1), 1–30.

Gu, Z., Gu, L., Eils, R., Schlesner, M., & Brors, B. (2014). Circlize implements and enhances circular visualization in R. Bioinformatics, 30(19), 2811–2812.

Hämälä, T., Wafula, E. K., Guiltinan, M. J., Ralph, P. E., dePamphilis, C. W., & Tiffin, P. (2021). Genomic structural variants constrain and facilitate adaptation in natural populations of Theobroma cacao, the chocolate tree. Proceedings of the National Academy of Sciences, 118(35).

Hardie, D. C., & Hebert, P. D. (2003). The nucleotypic effects of cellular DNA content in cartilaginous and ray-finned fishes. Genome, 46(4), 683–706.

Harris, R. S. (2007). Improved pairwise alignment of genomic DNA. The Pennsylvania State University.

Hejase, H. A., Salman-Minkov, A., Campagna, L., Hubisz, M. J., Lovette, I. J., Gronau, I., & Siepel, A. (2020). Genomic islands of differentiation in a rapid avian radiation have been driven by recent selective sweeps. Proceedings of the National Academy of Sciences, 117(48), 30554. doi: 10.1073/pnas.2015987117

Heller, D., & Vingron, M. (2019). SVIM: structural variant identification using mapped long reads. Bioinformatics, 35(17), 2907–2915.

Heller, D., & Vingron, M. (2020). SVIM-asm: Structural variant detection from haploid and diploid genome assemblies. Bioinformatics, 36(22–23), 5519–5521.

Henderson, E. C., & Brelsford, A. (2020). Genomic differentiation across the speciation continuum in three hummingbird species pairs. BMC Evolutionary Biology, 20(1), 1–11.

Hickey, G., Heller, D., Monlong, J., Sibbesen, J. A., Sirén, J., Eizenga, J., … Paten, B. (2020). Genotyping structural variants in pangenome graphs using the vg toolkit. Genome Biology, 21(1), 1–17.

Ho, E. K. H., Bellis, E. S., Calkins, J., Adrion, J. R., Latta IV, L. C., & Schaack, S. (2021). Engines of change: Transposable element mutation rates are high and variable within Daphnia magna. PLOS Genetics, 17(11), e1009827. doi: 10.1371/journal.pgen.1009827

Ho, Urban, A. E., & Mills, R. E. (2019). Structural variation in the sequencing era. Nature Reviews Genetics, 21(3), 171–189.

Hoffmann, A. A., & Rieseberg, L. H. (2008). Revisiting the impact of inversions in evolution: From population genetic markers to drivers of adaptive shifts and speciation? Annual Review of Ecology, Evolution, and Systematics, 39, 21–42.

Huddleston, J., Ranade, S., Malig, M., Antonacci, F., Chaisson, M., Hon, L., … Dennis, M. Y. (2014). Reconstructing complex regions of genomes using long-read sequencing technology. Genome Research, gr–168450.

Jacobsen, M. W., Hansen, M. M., Orlando, L., Bekkevold, D., Bernatchez, L., Willerslev, E., & Gilbert, M. T. P. (2012). Mitogenome sequencing reveals shallow evolutionary histories and recent divergence time between morphologically and ecologically distinct European whitefish (Coregonus spp.). Molecular Ecology, 21(11), 2727–2742.

Jain, C., Rhie, A., Zhang, H., Chu, C., Walenz, B. P., Koren, S., & Phillippy, A. M. (2020). Weighted minimizer sampling improves long read mapping. Bioinformatics, 36(Supplement_1), i111–i118.

Jiggins, C. D. (2019). Can genomics shed light on the origin of species? PLOS Biology, 17(8), e3000394. doi: 10.1371/journal.pbio.3000394

Kirkpatrick, M., & Barton, N. (2006). Chromosome inversions, local adaptation and speciation. Genetics, 173(1), 419–434.

Kirsche, M., Prabhu, G., Sherman, R., Ni, B., Aganezov, S., & Schatz, M. C. (2021). Jasmine: Population-scale structural variant comparison and analysis. BioRxiv.

Kolmogorov, M., Yuan, J., Lin, Y., & Pevzner, P. A. (2019). Assembly of long, error-prone reads using repeat graphs. Nature Biotechnology, 37(5), 540–546.

Korneliussen, T. S., Albrechtsen, A., & Nielsen, R. (2014). ANGSD: analysis of next generation sequencing data. BMC Bioinformatics, 15(1), 356.

Landis, J. B., Soltis, D. E., Li, Z., Marx, H. E., Barker, M. S., Tank, D. C., & Soltis, P. S. (2018). Impact of whole-genome duplication events on diversification rates in angiosperms. American Journal of Botany, 105(3), 348–363.

Landry, L., Vincent, W., & Bernatchez, L. (2007). Parallel evolution of lake whitefish dwarf ecotypes in association with limnological features of their adaptive landscape. Journal of Evolutionary Biology, 20(3), 971–984.

Laporte, M, Le Luyer, J., Rougeux, C., Dion-Côté, A.-M., Krick, M., & Bernatchez, L. (2019). DNA methylation reprogramming, TE derepression, and postzygotic isolation of nascent animal species. Science Advances, 5(10), eaaw1644.

Laporte, Martin, Dalziel, A. C., Martin, N., & Bernatchez, L. (2016). Adaptation and acclimation of traits associated with swimming capacity in Lake Whitefish (Coregonus clupeaformis) ecotypes. BMC Evolutionary Biology, 16(1), 1–13.

Laporte, Martin, Rogers, S. M., Dion-Côté, A.-M., Normandeau, E., Gagnaire, P.-A., Dalziel, A. C., … Bernatchez, L. (2015). RAD-QTL mapping reveals both genome-level parallelism and different genetic architecture underlying the evolution of body shape in lake whitefish (Coregonus clupeaformis) species pairs. G3: Genes, Genomes, Genetics, 5(7), 1481–1491.

Layer, R. M., Chiang, C., Quinlan, A. R., & Hall, I. M. (2014). LUMPY: a probabilistic framework for structural variant discovery. Genome Biology, 15(6), R84.

Legendre, P., & Legendre, L. F. (2012). Numerical ecology (Vol. 24). Elsevier.

Lemay, M.-A., Sibbesen, J. A., Torkamaneh, D., Hamel, J., Levesque, R. C., & Belzile, F. (2021). Combined use of Oxford Nanopore and Illumina sequencing yields insights into soybean structural variation biology. BioRxiv.

Li, H. (2018). Minimap2: Pairwise alignment for nucleotide sequences. Bioinformatics, 34(18), 3094–3100.

Li, H., & Durbin, R. (2009). Fast and accurate short read alignment with Burrows–Wheeler transform. Bioinformatics, 25(14), 1754–1760.

Li, H., Feng, X., & Chu, C. (2020). The design and construction of reference pangenome graphs with minigraph. Genome Biology, 21(1), 1–19.

Li, H., Handsaker, B., Wysoker, A., Fennell, T., Ruan, J., Homer, N., … Durbin, R. (2009). The sequence alignment/map format and SAMtools. Bioinformatics, 25(16), 2078–2079.

Lien, S., Koop, B. F., Sandve, S. R., Miller, J. R., Kent, M. P., Nome, T., … Zimin, A. (2016). The Atlantic salmon genome provides insights into rediploidization. Nature, 533(7602), 200–205.

Limborg, M. T., McKinney, G. J., Seeb, L. W., & Seeb, J. E. (2016). Recombination patterns reveal information about centromere location on linkage maps. Molecular Ecology Resources, 16(3), 655–661.

Lockwood, S. F., Seavey, B. T., Dillinger Jr, R. E., & Bickham, J. W. (1991). Variation in DNA content among age classes of broad whitefish (Coregonus nasus) from the Sagavanirktok River delta. Canadian Journal of Zoology, 69(5), 1335–1338.

Logsdon, G. A., Vollger, M. R., & Eichler, E. E. (2020). Long-read human genome sequencing and its applications. Nature Reviews Genetics, 21(10), 597–614.

Lou, R. N., Jacobs, A., Wilder, A., & Therkildsen, N. O. (2020). A beginner’s guide to low-coverage whole genome sequencing for population genomics.

Lu, G., & Bernatchez, L. (1999). Correlated trophic specialization and genetic divergence in sympatric lake whitefish ecotypes (Coregonus clupeaformis): Support for the ecological speciation hypothesis. Evolution, 53(5), 1491–1505.

Macqueen, D. J., & Johnston, I. A. (2014). A well-constrained estimate for the timing of the salmonid whole genome duplication reveals major decoupling from species diversification. Proceedings of the Royal Society B: Biological Sciences, 281(1778), 20132881.

Mahmoud, M., Gobet, N., Cruz-Dávalos, D. I., Mounier, N., Dessimoz, C., & Sedlazeck, F. J. (2019). Structural variant calling: The long and the short of it. Genome Biology, 20(1), 246.

Marçais, G., Delcher, A. L., Phillippy, A. M., Coston, R., Salzberg, S. L., & Zimin, A. (2018). MUMmer4: A fast and versatile genome alignment system. PLoS Computational Biology, 14(1), 1005944.

Marques, D. A., Lucek, K., Meier, J. I., Mwaiko, S., Wagner, C. E., Excoffier, L., & Seehausen, O. (2016). Genomics of rapid incipient speciation in sympatric threespine stickleback. PLoS Genetics, 12(2), e1005887.

Martin, S. H., Davey, J. W., Salazar, C., & Jiggins, C. D. (2019). Recombination rate variation shapes barriers to introgression across butterfly genomes. PLoS Biology, 17(2), e2006288.

McKenna, A., Hanna, M., Banks, E., Sivachenko, A., Cibulskis, K., Kernytsky, A., … Daly, M. (2010). The Genome Analysis Toolkit: A MapReduce framework for analyzing next-generation DNA sequencing data. Genome Research, 20(9), 1297–1303.

Meier, J. I., Marques, D. A., Wagner, C. E., Excoffier, L., & Seehausen, O. (2018). Genomics of parallel ecological speciation in Lake Victoria cichlids. Molecular Biology and Evolution, 35(6), 1489–1506.

Meisner, J., & Albrechtsen, A. (2018). Inferring population structure and admixture proportions in low-depth NGS data. Genetics, 210(2), 719–731.

Mérot, C., Oomen, R. A., Tigano, A., & Wellenreuther, M. (2020). A roadmap for understanding the evolutionary significance of structural genomic variation. Trends in Ecology & Evolution, 35(7), 561–572.

Morales, H. E., Faria, R., Johannesson, K., Larsson, T., Panova, M., Westram, A. M., & Butlin, R. K. (2019). Genomic architecture of parallel ecological divergence: Beyond a single environmental contrast. Science Advances, 5(12), eaav9963.

Nadeau, N. J., & Kawakami, T. (2019). Population Genomics of Speciation and Admixture. In O. P. Rajora (Ed.), Population Genomics: Concepts, Approaches and Applications (pp. 613–653). Cham: Springer International Publishing. doi: 10.1007/13836_2018_24

Noor, M. A., Grams, K. L., Bertucci, L. A., Almendarez, Y., Reiland, J., & Smith, K. R. (2001). The genetics of reproductive isolation and the potential for gene exchange between Drosophila pseudoobscura and D. persimilis via backcross hybrid males. Evolution, 55(3), 512–521.

Ou, S., & Jiang, N. (2019). LTR_FINDER_parallel: Parallelization of LTR_FINDER enabling rapid identification of long terminal repeat retrotransposons. Mobile DNA, 10(1), 1–3.

Phillips, R. B., Reed, K. M., & Ráb, P. (1996). Revised karyotypes and chromosome banding of coregonid fishes from the Laurentian Great Lakes. Canadian Journal of Zoology, 74(2), 323–329.

Rastas, P. (2017). Lep-MAP3: Robust linkage mapping even for low-coverage whole genome sequencing data. Bioinformatics, 33(23), 3726–3732. doi: 10.1093/bioinformatics/btx494

Rausch, T., Zichner, T., Schlattl, A., Stütz, A. M., Benes, V., & Korbel, J. O. (2012). DELLY: structural variant discovery by integrated paired-end and split-read analysis. Bioinformatics, 28(18), i333–i339.

Ravinet, M., Faria, R., Butlin, R., Galindo, J., Bierne, N., Rafajlović, M., … Westram, A. (2017). Interpreting the genomic landscape of speciation: A road map for finding barriers to gene flow. Journal of Evolutionary Biology, 30(8), 1450–1477.

Renaut, S., & Bernatchez, L. (2011). Transcriptome-wide signature of hybrid breakdown associated with intrinsic reproductive isolation in lake whitefish species pairs (Coregonus spp. Salmonidae). Heredity, 106(6), 1003–1011.

Renaut, S., Nolte, A., & Bernatchez, L. (2009). Transcriptomic investigation of post-zygotic isolation in lake whitefish (Coregonus clupeaformis). Mol. Biol. Evol, 26, 925–936.

Rogers, & Bernatchez, L. (2007). The genetic architecture of ecological speciation and the association with signatures of selection in natural lake whitefish (Coregonus sp. Salmonidae) species pairs. Molecular Biology and Evolution, 24(6), 1423–1438.

Rogers, S., & Bernatchez, L. (2006). The genetic basis of intrinsic and extrinsic post-zygotic reproductive isolation jointly promoting speciation in the lake whitefish species complex (Coregonus clupeaformis). Journal of Evolutionary Biology, 19(6), 1979–1994.

Rogers, S. M., Gagnon, V., & Bernatchez, L. (2002). Genetically based phenotype-environment association for swimming behavior in lake whitefish ecotypes (Coregonus clupeaformis Mitchill). Evolution, 56(11), 2322–2329.

Rogers, S. M., Isabel, N., & Bernatchez, L. (2007). Linkage maps of the dwarf and normal lake whitefish (Coregonus clupeaformis) species complex and their hybrids reveal the genetic architecture of population divergence. Genetics, 175(1), 375–398.

Rougeux, Bernatchez, L., & Gagnaire, P.-A. (2017). Modeling the multiple facets of speciation-with-gene-flow toward inferring the divergence history of lake whitefish species pairs (Coregonus clupeaformis). Genome Biology and Evolution, 9(8), 2057–2074.

Rougeux, Gagnaire, P., Praebel, K., Seehausen, O., & Bernatchez, L. (2019). Polygenic selection drives the evolution of convergent transcriptomic landscapes across continents within a Nearctic sister species complex. Molecular Ecology, 28(19), 4388–4403.

Scannell, D. R., Byrne, K. P., Gordon, J. L., Wong, S., & Wolfe, K. H. (2006). Multiple rounds of speciation associated with reciprocal gene loss in polyploid yeasts. Nature, 440(7082), 341–345.

Sedlazeck, F. J., Rescheneder, P., Smolka, M., Fang, H., Nattestad, M., Von Haeseler, A., & Schatz, M. C. (2018). Accurate detection of complex structural variations using single-molecule sequencing. Nature Methods, 15(6), 461–468.

Seehausen, O., Butlin, R. K., Keller, I., Wagner, C. E., Boughman, J. W., Hohenlohe, P. A., … Brännström, Å. (2014). Genomics and the origin of species. Nature Reviews Genetics, 15(3), 176–192.

Serrato-Capuchina, A., & Matute, D. R. (2018). The role of transposable elements in speciation. Genes, 9(5), 254.

Sirén, J., Monlong, J., Chang, X., Novak, A. M., Eizenga, J. M., Markello, C., … Carroll, A. (2020). Genotyping common, large structural variations in 5,202 genomes using pangenomes, the Giraffe mapper, and the vg toolkit. BioRxiv.

Smith, S. R., Normandeau, E., Djambazian, H., Nawarathna, P. M., Berube, P., Muir, A. M., … Luikart, G. (2021). A chromosome-anchored genome assembly for Lake Trout (Salvelinus namaycush). Molecular Ecology Resources.

Soderlund, C., Bomhoff, M., & Nelson, W. M. (2011). SyMAP v3. 4: A turnkey synteny system with application to plant genomes. Nucleic Acids Research, 39(10), e68–e68.

Stevison, L. S., & McGaugh, S. E. (2020). It’s time to stop sweeping recombination rate under the genome scan rug. Molecular Ecology, 29(22), 4249–4253. doi: 10.1111/mec.15690

Sutherland, B. J. G., Gosselin, T., Normandeau, E., Lamothe, M., Isabel, N., Audet, C., & Bernatchez, L. (2016). Salmonid Chromosome Evolution as Revealed by a Novel Method for Comparing RADseq Linkage Maps. Genome Biology and Evolution, 8(12), 3600–3617. doi: 10.1093/gbe/evw262

Tham, C. Y., Tirado-Magallanes, R., Goh, Y., Fullwood, M. J., Koh, B. T., Wang, W., … Tenen, D. G. (2020). NanoVar: Accurate characterization of patients’ genomic structural variants using low-depth nanopore sequencing. Genome Biology, 21(1), 1–15.

Ting, C.-T., Tsaur, S.-C., Sun, S., Browne, W. E., Chen, Y.-C., Patel, N. H., & Wu, C.-I. (2004). Gene duplication and speciation in Drosophila: Evidence from the Odysseus locus. Proceedings of the National Academy of Sciences, 101(33), 12232–12235.

Todesco, M., Owens, G. L., Bercovich, N., Légaré, J.-S., Soudi, S., Burge, D. O., … Imerovski, I. (2020). Massive haplotypes underlie ecotypic differentiation in sunflowers. Nature, 584(7822), 602–607.

Walker, B. J., Abeel, T., Shea, T., Priest, M., Abouelliel, A., Sakthikumar, S., … Young, S. K. (2014). Pilon: An integrated tool for comprehensive microbial variant detection and genome assembly improvement. PloS One, 9(11), e112963.

Waples, R. K., Seeb, L. W., & Seeb, J. E. (2016). Linkage mapping with paralogs exposes regions of residual tetrasomic inheritance in chum salmon (Oncorhynchus keta). Molecular Ecology Resources, 16(1), 17–28.

Weissensteiner, M. H., Bunikis, I., Catalán, A., Francoijs, K.-J., Knief, U., Heim, W., … Suh, A. (2020). Discovery and population genomics of structural variation in a songbird genus. Nature Communications, 11(1), 1–11.

Wellenreuther, M., & Bernatchez, L. (2018). Eco-evolutionary genomics of chromosomal inversions. Trends in Ecology & Evolution, 33(6), 427–440.

Wolf, J. B., & Ellegren, H. (2017). Making sense of genomic islands of differentiation in light of speciation. Nature Reviews Genetics, 18(2), 87–100.

