## Supplementary materials for "Genome assembly, structural variants, and genetic differentiation between Lake Whitefish young species pairs (*Coregonus* sp.) with long and short reads"

Table S1: Statistics for the genome assemblies before anchoring into chromosomes.

| Whitefish genome assemblies | total length [bp] | number of contigs | longest contig [bp] | L50 [bp] | N50 [n] | L90 [bp] | N90 [n] |
|---|---|---|---|---|---|---|---|
| Normal Final assembly | 2682618941 | 7076 | 43030526 | 6096834 | 101 | 308820 | 870 |
| Normal Whitefish_Flye08K | 2712325911 | 8777 | 24349152 | 3219133 | 201 | 168561 | 1863 |
| Normal Whitefish_Flye10K | 2772957034 | 8762 | 25401145 | 2368812 | 257 | 166291 | 2143 |
| Normal Whitefish_Flye15K | 2785624403 | 9475 | 11362423 | 970569 | 707 | 131193 | 3746 |
| Dwarf Whitefish Flye10k | 2764848066 | 8433 | 21574413 | 2160360 | 274 | 159968 | 2272 |
| Dwarf Whitefish Flye08k | 2780381566 | 10128 | 21923556 | 2153278 | 291 | | |
| Dwarf Whitefish Flye12k | 2807411528 | 9494 | 14914173 | 1671611 | 377 | | |

### Table S2: Correspondence between chromosomes in Coregonids

Correspondence between the reference genomes of *Coregonus clupeaformis* and *C. lavaretus* and linkage groups in genetic maps of *Coregonus clupeaformis, C. artedii,* and *C. lavaretus*.

| Chromosome name in *Coregonus clupeaformis* genome (this study) | Chromosome name in *Coregonus lavaretus* genome (DeKayne et al, 2020) | Linkage group in *Coregonus lavaretus* genetic map (DeKayne & Feulner, 2018) | Linkage group in *Coregonus artedi* genetic map (Blumstein et al, 2020) | Linkage group in *Coregonus clupeaformis* previous genetic map (Gagnaire et al, 2013) | Linkage group in *Coregonus clupeaformis* new genetic map (this study) |
|---|---|---|---|---|---|
| Chr01 | WFS01 | 1 | 6 | 5 | 4 |
| Chr02 | WFS02 | 29 | 30 | 16 | 30 |
| Chr03 | WFS03 | 30 | 21 | 29 | 31 |
| Chr04 | WFS04 | 3 (+36) | 7 | 2 (+3)? | 34 |
| Chr05 | WFS05 | 16 (+33) | 23 | 28 | 3 |
| Chr06 | WFS06 | 33 (+16) | 14 | 35 | 18 |
| Chr07 | WFS07 | 13 (+34) | 5 | 12 | 15 |
| Chr08 | WFS08 | 9 | 4 | 24 | 2 |
| Chr09 | WFS09 | 6 | 20 | 13 | 14 |
| Chr10 | WFS10 | 5 | 24 | 38 | 5 |
| Chr11 | WFS11 | 15 | 11 | 11 | 11 |
| Chr12 | WFS12 | 12 | 22 | 21 | 16 |
| Chr13 | WFS13 | 7 | 19 | 8 | 19 |
| Chr14 | WFS14 | 22 | 26 | 26 | 1 |
| Chr15 | WFS15 | 2 | 1 | 4 | 12 |
| Chr16 | WFS16 | 25 | 17 | 34 | 21 |
| Chr17 | WFS17 | 20 | 8 | 10 | 10 |
| Chr18 | WFS18 | 31 | 32 | 37 | 25 |
| Chr19 | WFS19 | 4 | 13 | 30 | 6 |
| Chr20 | WFS20 | 8 | 3 | 6 | 7 |
| Chr21 | WFS21 | 19 | 25 | 25 | 24 |
| Chr22 | WFS22 | 38+39 | NA | 7 | 36 |
| Chr23 | WFS23 | 21 | 28 | 32 | 17 |
| Chr24 | WFS24 | 28 | 16 | 14 | 20 |
| Chr25 | WFS25 | 10 | 31 | 22 | 39 |
| Chr26 | WFS26 | 17 | 27 | 36 | 13 |
| Chr27 | WFS27 | 11 | 38 | 33 | 8 |
| Chr28 | WFS28 | 36 (+20) | 2 | 1 | 29 |
| Chr29 | WFS29 | 18 | 10 | 31 | 9 |
| Chr30 | WFS30 | 14 | 33 | 27 | 28 |
| Chr31 | WFS31 | 35 | 18 | 15 | 26 |
| Chr32 | WFS32 | 37 (+2) | 12 | 4 | 35 |
| Chr33 | WFS33 | 23 | 37 | 17 | 27 |
| Chr34 | WFS34 | 26 | 36 | 39 | 22 |
| Chr35 | WFS35 | 34 (+13) | 9 | NA/18? | 23 |
| Chr36 | WFS36 | 24 | 29 | 40 | 38 |
| Chr37 | WFS37 | 27 (+1) | 15 | 5 | 32 |
| Chr38 | WFS38 | 8 | 34 | 6 | 37 |
| Chr39 | WFS39 | 32 | 35 | 23 | 33 |
| Chr40 | NA | 3 (+36) | 7 | 2+3? | 40 |

Table S3: Coordinates of putatively collapsed duplicated regions in the reference assembly of the Normal Lake Whitefish

| | | |
|---|---|---|
| Chr01 | 18000000 | 20500000 |
| Chr01 | 103700000 | 121031747 |
| Chr03 | 43640000 | 45080000 |
| Chr04 | 32010000 | 34920000 |
| Chr07 | 11410000 | 12870000 |
| Chr07 | 81600000 | 82980000 |
| Chr10 | 62190000 | 66060000 |
| Chr13 | 1 | 3020000 |
| Chr15 | 73710000 | 75020000 |
| Chr18 | 26260000 | 27550000 |
| Chr20 | 1800000 | 8400000 |
| Chr22 | 1 | 7216770 |
| Chr24 | 1 | 8220000 |
| Chr28 | 20000000 | 49420000 |
| Chr30 | 56520000 | 64630000 |
| Chr32 | 1 | 11602415 |
| Chr35 | 20840000 | 23000000 |
| Chr38 | 1 | 8900000 |
| Chr39 | 1 | 7110000 |

Table S4: Proportion of interspersed repeats in the Normal genome and in the sequences of the structural variants

| | Normal reference genome | | | Sequences of SVs | | |
|---|---|---|---|---|---|---|
| | number of sequences | length of sequences | % of the length | number of sequences | length of sequences | % of the length |
| Unmasked | | 1008956356 | 37.61% | | 49088355 | 21.63% |
| Retroelements | 1635898 | 694575878 | 25.89% | 162140 | 111518063 | 49.13% |
| DNA-transposons | 1587932 | 629778518 | 23.48% | 96759 | 42683508 | 18.81% |
| Rolling-circles | 8595 | 1329769 | 0.05% | 745 | 139058 | 0.06% |
| Unclassified TE | 1505716 | 246978169 | 9.21% | 64010 | 9506168 | 4.19% |
| Small RNA | 121294 | 19679506 | 0.73% | 8423 | 1613074 | 0.71% |
| Satellites | 7644 | 1993471 | 0.07% | 454 | 86506 | 0.04% |
| Simple repeats | 727239 | 68691185 | 2.56% | 110507 | 11415712 | 5.03% |
| Low complexity | 84004 | 10696989 | 0.40% | 5018 | 920708 | 0.41% |

Table S5: Breakdown of the proportion of transposable elements for the main families in the Normal genome and in the sequences of the structural variants

| | Normal reference genome | | | Sequences of SVs | | |
|---|---|---|---|---|---|---|
| | number of sequences | length of sequences | % of the length | number of sequences | length of sequences | % of the length |
| Retroelements/SINEs | 165982 | 19369894 | 0.72% | 11103 | 1858884 | 0.82% |
| Retroelements/Penelope | 16711 | 3023382 | 0.11% | 573 | 89414 | 0.04% |
| Retroelements/LINEs | 752039 | 336040143 | 12.53% | 48725 | 28036418 | 12.35% |
| Retroelements/CRE/SLACS | 0 | 0 | 0% | 0 | 0 | 0% |
| Retroelements/L2/CR1/Rex | 527901 | 235787647 | 8.79% | 35376 | 21505242 | 9.47% |
| Retroelements/R1/LOA/Jockey | 22192 | 7475316 | 0.28% | 1042 | 324057 | 0.14% |
| Retroelements/R2/R4/NeSL | 816 | 538660 | 0.02% | 56 | 10508 | 0% |
| Retroelements/RTE/Bov-B | 67311 | 39504265 | 1.47% | 5650 | 2598236 | 1.14% |
| Retroelements/L1/CIN4 | 27755 | 11837512 | 0.44% | 1462 | 1461716 | 0.64% |
| Retroelements/LTR | 717877 | 339165841 | 12.64% | 102312 | 81622761 | 35.96% |
| Retroelements/BEL/Pao | 1667 | 3089456 | 0.12% | 282 | 693587 | 0.31% |
| Retroelements/Ty1/Copia | 6651 | 1186106 | 0.04% | 486 | 198262 | 0.09% |
| Retroelements/Gypsy/DIRS1 | 215716 | 177453473 | 6.61% | 30682 | 51256082 | 22.58% |
| Retroelements/Retroviral | 94119 | 37728202 | 1.41% | 11140 | 5625760 | 2.48% |
| DNA-transposons/hobo-Activator | 215774 | 54227588 | 2.02% | 13741 | 5110005 | 2.25% |
| DNA-transposons/Tc1-IS630-Pogo | 1156068 | 538733044 | 20.08% | 67533 | 33021547 | 14.55% |
| DNA-transposons/En-Spm | 0 | 0 | 0% | 0 | 0 | 0% |
| DNA-transposons/MuDR-IS905 | 0 | 0 | 0% | 0 | 0 | 0% |
| DNA-transposons/PiggyBac | 10077 | 3142994 | 0.12% | 546 | 253304 | 0.11% |
| DNA-transposons/Tourist/Harbinger | 11323 | 3509455 | 0.13% | 825 | 547015 | 0.24% |
| DNA-transposons/Other | 719 | 201371 | 0.01% | 25 | 4403 | 0% |

Table S6: Portion of the genome masked by different families of TEs

| | size in Dwarf | % of the Dwarf Genome | size in Normal | % of the Normal genome |
|---|---|---|---|---|
| 5S-Deu-L2 | 4580718 | 0.17% | 4139986 | 0.15% |
| CMC-EnSpm | 8615241 | 0.31% | 8447378 | 0.31% |
| Copia | 1103875 | 0.04% | 1057340 | 0.04% |
| Crypton-A | 812641 | 0.03% | 799915 | 0.03% |
| Crypton-V | 4066459 | 0.15% | 4562557 | 0.17% |
| DIRS | 6594109 | 0.24% | 6364078 | 0.23% |
| DNA | 8085117 | 0.29% | 7730840 | 0.28% |
| ERV | 1787503 | 0.06% | 1749186 | 0.06% |
| ERV1 | 37428917 | 1.35% | 37122674 | 1.36% |
| Ginger-1 | 147473 | 0.01% | 147412 | 0.01% |
| Gypsy | 176462088 | 6.38% | 171307131 | 6.26% |
| hAT | 3200603 | 0.12% | 3117699 | 0.11% |
| hAT-Ac | 17428675 | 0.63% | 17411111 | 0.64% |
| hAT-Blackjack | 1192194 | 0.04% | 1175813 | 0.04% |
| hAT-Charlie | 16524837 | 0.60% | 16193170 | 0.59% |
| hAT-hAT5 | 504449 | 0.02% | 486359 | 0.02% |
| hAT-hAT6 | 678618 | 0.02% | 678122 | 0.02% |
| hAT-Tip100 | 17333573 | 0.63% | 16047021 | 0.59% |
| Helitron | 1283837 | 0.05% | 1252969 | 0.05% |
| I | 8078027 | 0.29% | 7744404 | 0.28% |
| IS3EU | 793720 | 0.03% | 720151 | 0.03% |
| Kolobok-E | 650086 | 0.02% | 608333 | 0.02% |
| Kolobok-T2 | 1096387 | 0.04% | 1089204 | 0.04% |
| L1 | 11284387 | 0.41% | 12025585 | 0.44% |
| L1-Tx1 | 39839302 | 1.44% | 38773224 | 1.42% |
| L2 | 142551018 | 5.16% | 138666129 | 5.07% |
| LINE | 1046032 | 0.04% | 1050364 | 0.04% |
| LTR | 29955 | 0.00% | 29014 | 0.00% |
| Maverick | 1878546 | 0.07% | 1795159 | 0.07% |
| Merlin | 175903 | 0.01% | 179830 | 0.01% |
| Ngaro | 230312 | 0.01% | 217572 | 0.01% |
| P | 206746 | 0.01% | 201657 | 0.01% |
| Pao | 3125866 | 0.11% | 3069278 | 0.11% |
| Penelope | 2681214 | 0.10% | 2856849 | 0.10% |
| PIF | 137083 | 0.00% | 128569 | 0.00% |
| PIF-Harbinger | 3832933 | 0.14% | 3529906 | 0.13% |
| PIF-ISL2EU | 2010155 | 0.07% | 2022879 | 0.07% |
| PiggyBac | 3058264 | 0.11% | 3141712 | 0.11% |
| R2-NeSL | 431679 | 0.02% | 537377 | 0.02% |
| Rex-Babar | 97757592 | 3.54% | 96050620 | 3.51% |
| RTE-BovB | 154480 | 0.01% | 150975 | 0.01% |
| RTE-X | 41541679 | 1.50% | 56048671 | 2.05% |
| SINE | 2013974 | 0.07% | 1956691 | 0.07% |
| SINE? | 8581657 | 0.31% | 9841653 | 0.36% |
| Sola-1 | 283780 | 0.01% | 288855 | 0.01% |
| Sola-2 | 763024 | 0.03% | 737761 | 0.03% |
| TcMar | 456471 | 0.02% | 410609 | 0.02% |
| TcMar-Fot1 | 2045407 | 0.07% | 2882523 | 0.11% |
| TcMar-ISRm11 | 8253620 | 0.30% | 7751729 | 0.28% |
| TcMar-Tc1 | 543460096 | 19.66% | 533614040 | 19.50% |
| TcMar-Tc2 | 1031109 | 0.04% | 1071410 | 0.04% |
| TcMar-Tigger | 991499 | 0.04% | 932645 | 0.03% |
| tRNA-Core-RTE | 2423485 | 0.09% | 2549640 | 0.09% |
| tRNA-Deu-RTE | 2727426 | 0.10% | 2667158 | 0.10% |
| Unknown | 379770302 | 13.74% | 387922260 | 14.18% |
| Zisupton | 580639 | 0.02% | 544959 | 0.02% |

Table S7: Enrichment in SVs associated with transposable elements in outliers of differentiation between Dwarf and Normal Whitefish.

| Element associated with the SV | Dataset for population-level analysis | | Outliers in Indian Lake | | | | | Outliers in Cliff Lake | | | | | Overlap of outlier SVs across lakes | | | | |
|---|---|---|---|---|---|---|---|---|---|---|---|---|---|---|---|---|---|
| | N | % | N | % | p | OR | q | N | % | p | OR | q | N | % | p | OR | q |
| DNA | 93 | 0.1% | 5 | 0.1% | 0.502 | 1.1 | 0.887 | 7 | 0.1% | 0.200 | 1.5 | 0.619 | - | - | 1.000 | 0.0 | 1.000 |
| DNA/CMC-EnSpm | 78 | 0.1% | 6 | 0.1% | 0.211 | 1.5 | 0.453 | 3 | 0.1% | 0.747 | 0.8 | 1.000 | - | - | 1.000 | 0.0 | 1.000 |
| DNA/Crypton-A | 7 | 0.0% | - | - | 1.000 | 0.0 | 1.000 | 1 | 0.0% | 0.323 | 2.9 | 0.762 | - | - | 1.000 | 0.0 | 1.000 |
| DNA/Crypton-V | 106 | 0.1% | 9 | 0.2% | 0.099 | 1.7 | 0.269 | 4 | 0.1% | 0.774 | 0.8 | 1.000 | 2 | 0.3% | 0.203 | 2.4 | 0.782 |
| DNA/hAT | 24 | 0.0% | 2 | 0.0% | 0.353 | 1.7 | 0.662 | 1 | 0.0% | 0.705 | 0.8 | 1.000 | - | - | 1.000 | 0.0 | 1.000 |
| DNA/hAT-Ac | 484 | 0.5% | **57** | **1.2%** | **0.000** | **2.4** | **0.000** | **37** | **0.8%** | **0.011** | **1.5** | **0.057** | **12** | **1.7%** | **0.001** | **3.2** | **0.009** |
| DNA/hAT-Blackjack | 15 | 0.0% | - | - | 1.000 | 0.0 | 1.000 | 2 | 0.0% | 0.193 | 2.7 | 0.619 | - | - | 1.000 | 0.0 | 1.000 |
| DNA/hAT-Charlie | 528 | 0.6% | **49** | **1.0%** | **0.000** | **1.9** | **0.001** | **50** | **1.1%** | **0.000** | **1.9** | **0.000** | 9 | 1.2% | 0.026 | 2.2 | 0.192 |
| DNA/hAT-hAT5 | 3 | 0.0% | - | - | 1.000 | 0.0 | 1.000 | - | - | 1.000 | 0.0 | 1.000 | - | - | 1.000 | 0.0 | 1.000 |
| DNA/hAT-hAT6 | 5 | 0.0% | - | - | 1.000 | 0.0 | 1.000 | - | - | 1.000 | 0.0 | 1.000 | - | - | 1.000 | 0.0 | 1.000 |
| DNA/hAT-Tip100 | 301 | 0.3% | **29** | **0.6%** | **0.001** | **1.9** | **0.007** | 19 | 0.4% | 0.193 | 1.3 | 0.619 | 2 | 0.3% | 0.678 | 0.9 | 1.000 |
| DNA/IS3EU | 3 | 0.0% | 1 | 0.0% | 0.177 | 6.7 | 0.426 | - | - | 1.000 | 0.0 | 1.000 | - | - | 1.000 | 0.0 | 1.000 |
| DNA/Kolobok-E | 8 | 0.0% | - | - | 1.000 | 0.0 | 1.000 | 1 | 0.0% | 0.355 | 2.5 | 0.762 | - | - | 1.000 | 0.0 | 1.000 |
| DNA/Kolobok-T2 | 21 | 0.0% | 1 | 0.0% | 0.658 | 1.0 | 1.000 | 1 | 0.0% | 0.658 | 1.0 | 1.000 | - | - | 1.000 | 0.0 | 1.000 |
| DNA/Maverick | 60 | 0.1% | 5 | 0.1% | 0.197 | 1.7 | 0.453 | 2 | 0.0% | 0.801 | 0.7 | 1.000 | 1 | 0.1% | 0.376 | 2.1 | 1.000 |
| DNA/PIF | 1 | 0.0% | - | - | 1.000 | 0.0 | 1.000 | - | - | 1.000 | 0.0 | 1.000 | - | - | 1.000 | 0.0 | 1.000 |
| DNA/PIF-Harbinger | 57 | 0.1% | 2 | 0.0% | 0.778 | 0.7 | 1.000 | 2 | 0.0% | 0.778 | 0.7 | 1.000 | - | - | 1.000 | 0.0 | 1.000 |
| DNA/PIF-ISL2EU | 69 | 0.1% | 5 | 0.1% | 0.277 | 1.4 | 0.554 | 4 | 0.1% | 0.461 | 1.2 | 0.937 | - | - | 1.000 | 0.0 | 1.000 |
| DNA/PiggyBac | 17 | 0.0% | 3 | 0.1% | 0.067 | 3.5 | 0.237 | 3 | 0.1% | 0.067 | 3.5 | 0.310 | 1 | 0.1% | 0.130 | 7.6 | 0.709 |

| | | | | | | | | | | | | | | | | | |
|---|---|---|---|---|---|---|---|---|---|---|---|---|---|---|---|---|---|
| DNA/Sola-1 | 3 | 0.0% | - | - | 1.000 | 0.0 | 1.000 | - | - | 1.000 | 0.0 | 1.000 | - | - | 1.000 | 0.0 | 1.000 |
| DNA/Sola-2 | 8 | 0.0% | 2 | 0.0% | 0.079 | 5.0 | 0.243 | 1 | 0.0% | 0.355 | 2.5 | 0.762 | - | - | 1.000 | 0.0 | 1.000 |
| DNA/TcMar | 1 | 0.0% | - | - | 1.000 | 0.0 | 1.000 | - | - | 1.000 | 0.0 | 1.000 | - | - | 1.000 | 0.0 | 1.000 |
| DNA/TcMar-Fot1 | 24 | 0.0% | 3 | 0.1% | 0.136 | 2.5 | 0.339 | 2 | 0.0% | 0.353 | 1.7 | 0.762 | - | - | 1.000 | 0.0 | 1.000 |
| DNA/TcMar-ISRm11 | 285 | 0.3% | 17 | 0.4% | 0.275 | 1.2 | 0.554 | 18 | 0.4% | 0.200 | 1.3 | 0.619 | - | - | 1.000 | 0.0 | 1.000 |
| DNA/TcMar-Tc1 | 11594 | 12.4% | **870** | **18.6%** | **0.000** | **1.5** | **0.000** | **876** | **18.7%** | **0.000** | **1.5** | **0.000** | **165** | **22.7%** | **0.000** | **1.8** | **0.000** |
| DNA/TcMar-Tc2 | 118 | 0.1% | 11 | 0.2% | 0.045 | 1.9 | 0.168 | 9 | 0.2% | 0.153 | 1.5 | 0.612 | 1 | 0.1% | 0.601 | 1.1 | 1.000 |
| DNA/TcMar-Tigger | 55 | 0.1% | 3 | 0.1% | 0.526 | 1.1 | 0.902 | 1 | 0.0% | 0.935 | 0.4 | 1.000 | - | - | 1.000 | 0.0 | 1.000 |
| DNA/Zisupton | 2 | 0.0% | - | - | 1.000 | 0.0 | 1.000 | - | - | 1.000 | 0.0 | 1.000 | - | - | 1.000 | 0.0 | 1.000 |
| LINE | 3 | 0.0% | - | - | 1.000 | 0.0 | 1.000 | - | - | 1.000 | 0.0 | 1.000 | - | - | 1.000 | 0.0 | 1.000 |
| LINE/I | 64 | 0.1% | **9** | **0.2%** | **0.008** | **2.8** | **0.036** | 5 | 0.1% | 0.232 | 1.6 | 0.632 | - | - | 1.000 | 0.0 | 1.000 |
| LINE/L1 | 96 | 0.1% | **8** | **0.2%** | **0.124** | **1.7** | **0.322** | **14** | **0.3%** | **0.001** | **2.9** | **0.005** | 2 | 0.3% | 0.175 | 2.7 | 0.782 |
| LINE/L1-Tx1 | 519 | 0.6% | 34 | 0.7% | 0.081 | 1.3 | 0.243 | 34 | 0.7% | 0.081 | 1.3 | 0.347 | 6 | 0.8% | 0.222 | 1.5 | 0.782 |
| LINE/L2 | 3076 | 3.3% | **231** | **4.9%** | **0.000** | **1.5** | **0.000** | **260** | **5.5%** | **0.000** | **1.7** | **0.000** | **37** | **5.1%** | **0.009** | **1.6** | **0.076** |
| LINE/Penelope | 24 | 0.0% | 1 | 0.0% | 0.705 | 0.8 | 1.000 | 1 | 0.0% | 0.705 | 0.8 | 1.000 | 1 | 0.1% | 0.176 | 5.4 | 0.782 |
| LINE/R2-NeSL | 7 | 0.0% | - | - | 1.000 | 0.0 | 1.000 | - | - | 1.000 | 0.0 | 1.000 | - | - | 1.000 | 0.0 | 1.000 |
| LINE/Rex-Babar | 1821 | 1.9% | **172** | **3.7%** | **0.000** | **1.9** | **0.000** | **157** | **3.3%** | **0.000** | **1.7** | **0.000** | 21 | 2.9% | 0.055 | 1.5 | 0.364 |
| LINE/RTE-BovB | 2 | 0.0% | 2 | 0.0% | 0.013 | 20.0 | 0.055 | - | - | 1.000 | 0.0 | 1.000 | - | - | 1.000 | 0.0 | 1.000 |
| LINE/RTE-X | 1113 | 1.2% | 28 | 0.6% | 1.000 | 0.5 | 1.000 | 39 | 0.8% | 0.991 | 0.7 | 1.000 | 3 | 0.4% | 0.991 | 0.3 | 1.000 |
| Low_complexity | 774 | 0.8% | 11 | 0.2% | 1.000 | 0.3 | 1.000 | 17 | 0.4% | 1.000 | 0.4 | 1.000 | 2 | 0.3% | 0.982 | 0.3 | 1.000 |
| LTR | 4 | 0.0% | - | - | 1.000 | 0.0 | 1.000 | 1 | 0.0% | 0.217 | 5.0 | 0.619 | - | - | 1.000 | 0.0 | 1.000 |
| LTR/Copia | 16 | 0.0% | 2 | 0.0% | 0.211 | 2.5 | 0.453 | 1 | 0.0% | 0.564 | 1.2 | 1.000 | - | - | 1.000 | 0.0 | 1.000 |
| LTR/DIRS | 91 | 0.1% | 10 | 0.2% | 0.022 | 2.2 | 0.090 | 5 | 0.1% | 0.484 | 1.1 | 0.937 | - | - | 1.000 | 0.0 | 1.000 |
| LTR/ERV | 21 | 0.0% | 1 | 0.0% | 0.658 | 1.0 | 1.000 | 1 | 0.0% | 0.658 | 1.0 | 1.000 | - | - | 1.000 | 0.0 | 1.000 |
| LTR/ERV1 | 594 | 0.6% | **54** | **1.2%** | **0.000** | **1.8** | **0.000** | **49** | **1.0%** | **0.001** | **1.6** | **0.006** | 7 | 1.0% | 0.186 | 1.5 | 0.782 |

| | | | | | | | | | | | | | | | | | |
|---|---|---|---|---|---|---|---|---|---|---|---|---|---|---|---|---|---|
| LTR/Gypsy | 5543 | 5.9% | **384** | **8.2%** | **0.000** | **1.4** | **0.000** | **481** | **10.3%** | **0.000** | **1.7** | **0.000** | **61** | **8.4%** | **0.007** | **1.4** | **0.073** |
| LTR/Ngaro | 2 | 0.0% | - | - | 1.000 | 0.0 | 1.000 | - | - | 1.000 | 0.0 | 1.000 | - | - | 1.000 | 0.0 | 1.000 |
| LTR/Pao | 46 | 0.1% | 5 | 0.1% | 0.095 | 2.2 | 0.269 | 2 | 0.0% | 0.673 | 0.9 | 1.000 | 1 | 0.1% | 0.305 | 2.8 | 0.962 |
| LTR/Unknown | 4374 | 4.7% | **334** | **7.1%** | **0.000** | **1.5** | **0.000** | **343** | **7.3%** | **0.000** | **1.6** | **0.000** | **51** | **7.0%** | **0.005** | **1.5** | **0.057** |
| no_TE | 30082 | 32.1% | 1494 | 31.9% | 0.594 | 1.0 | 0.963 | 1462 | 31.2% | 0.827 | 1.0 | 1.000 | 213 | 29.3% | 0.885 | 0.9 | 1.000 |
| RC/Helitron | 21 | 0.0% | - | - | 1.000 | 0.0 | 1.000 | - | - | 1.000 | 0.0 | 1.000 | - | - | 1.000 | 0.0 | 1.000 |
| rRNA | 12 | 0.0% | 1 | 0.0% | 0.470 | 1.7 | 0.854 | 1 | 0.0% | 0.470 | 1.7 | 0.937 | - | - | 1.000 | 0.0 | 1.000 |
| Satellite | 35 | 0.0% | 1 | 0.0% | 0.827 | 0.6 | 1.000 | 3 | 0.1% | 0.271 | 1.7 | 0.706 | 1 | 0.1% | 0.243 | 3.7 | 0.810 |
| Simple_repeat | 24142 | 25.8% | 232 | 4.9% | 1.000 | 0.2 | 1.000 | 181 | 3.9% | 1.000 | 0.1 | 1.000 | 22 | 3.0% | 1.000 | 0.1 | 1.000 |
| SINE | 22 | 0.0% | - | - | 1.000 | 0.0 | 1.000 | - | - | 1.000 | 0.0 | 1.000 | - | - | 1.000 | 0.0 | 1.000 |
| SINE/5S-Deu-L2 | 17 | 0.0% | 1 | 0.0% | 0.585 | 1.2 | 0.963 | - | - | 1.000 | 0.0 | 1.000 | - | - | 1.000 | 0.0 | 1.000 |
| SINE/tRNACore-RTE | 53 | 0.1% | 4 | 0.1% | 0.288 | 1.5 | 0.557 | 4 | 0.1% | 0.288 | 1.5 | 0.719 | 1 | 0.1% | 0.341 | 2.4 | 1.000 |
| SINE/tRNADeu-RTE | 30 | 0.0% | 4 | 0.1% | 0.077 | 2.7 | 0.243 | 3 | 0.1% | 0.207 | 2.0 | 0.619 | 1 | 0.1% | 0.213 | 4.3 | 0.782 |
| SINE? | 2132 | 2.3% | **198** | **4.2%** | **0.000** | **1.9** | **0.000** | **200** | **4.3%** | **0.000** | **1.9** | **0.000** | 37 | 5.1% | 0.000 | 2.2 | 0.000 |
| tRNA | 2291 | 2.4% | **158** | **3.4%** | **0.000** | **1.4** | **0.001** | **191** | **4.1%** | **0.000** | **1.7** | **0.000** | 24 | 3.3% | 0.096 | 1.4 | 0.575 |
| Unknown | 2776 | 3.0% | **230** | **4.9%** | **0.000** | **1.7** | **0.000** | **190** | **4.1%** | **0.000** | **1.4** | **0.000** | **43** | **5.9%** | **0.000** | **2.0** | **0.001** |

### Table S9: Coordinates of previously identified QTL in the new reference genome

Regions including markers identified as significantly associated with one of the phenotypic traits of interest by (Gagnaire, Normandeau, Pavey, & Bernatchez, 2013; Rogers, Isabel, & Bernatchez, 2007)

| CHR | start | stop | QTL |
|---|---|---|---|
| Chr02 | 24354669 | 24354670 | Maturity |
| Chr05 | 30243484 | 44560907 | Activity & Depth_selection |
| Chr05 | 47955978 | 47955979 | Gonadosomatic index |
| Chr06 | 10944482 | 11510663 | Growth rate |
| Chr07 | 13382789 | 13382790 | Directional change |
| Chr07 | 45142150 | 49941487 | Activity |
| Chr07 | 52999016 | 52999017 | Growth rate |
| Chr08 | 85507277 | 85507278 | Burst swimming & directional change |
| Chr10 | 53381623 | 60034589 | Depth selection |
| Chr12 | 51988913 | 52856337 | Depth selection |
| Chr13 | 10099482 | 63978158 | Gill raker |
| Chr13 | 46005064 | 47196556 | Depth selection |
| Chr15 | 23283752 | 56663130 | Gill raker |
| Chr17 | 164510 | 68091188 | Directional change |
| Chr20 | 27334774 | 27334775 | Directional change |
| Chr21 | 1967998 | 45849371 | sex |
| Chr23 | 11426032 | 54324241 | Directional change |
| Chr23 | 28790621 | 38959612 | Growth rate |
| Chr28 | 25985678 | 26432681 | Directional change |
| Chr29 | 2195646 | 6155549 | Gill raker |
| Chr31 | 8131059 | 12403428 | Activity |
| Chr33 | 38762999 | 41235318 | Gonadosomatic index |
| Chr34 | 30151992 | 35406463 | Depth selection |
| Chr35 | 21065802 | 31659917 | Directional change & Activity |
| Chr35 | 44842935 | 60738884 | Burst swimming |
| Chr39 | 11171630 | 30309175 | Depth selection |
| Chr40 | 1267440 | 11926388 | Maturity |

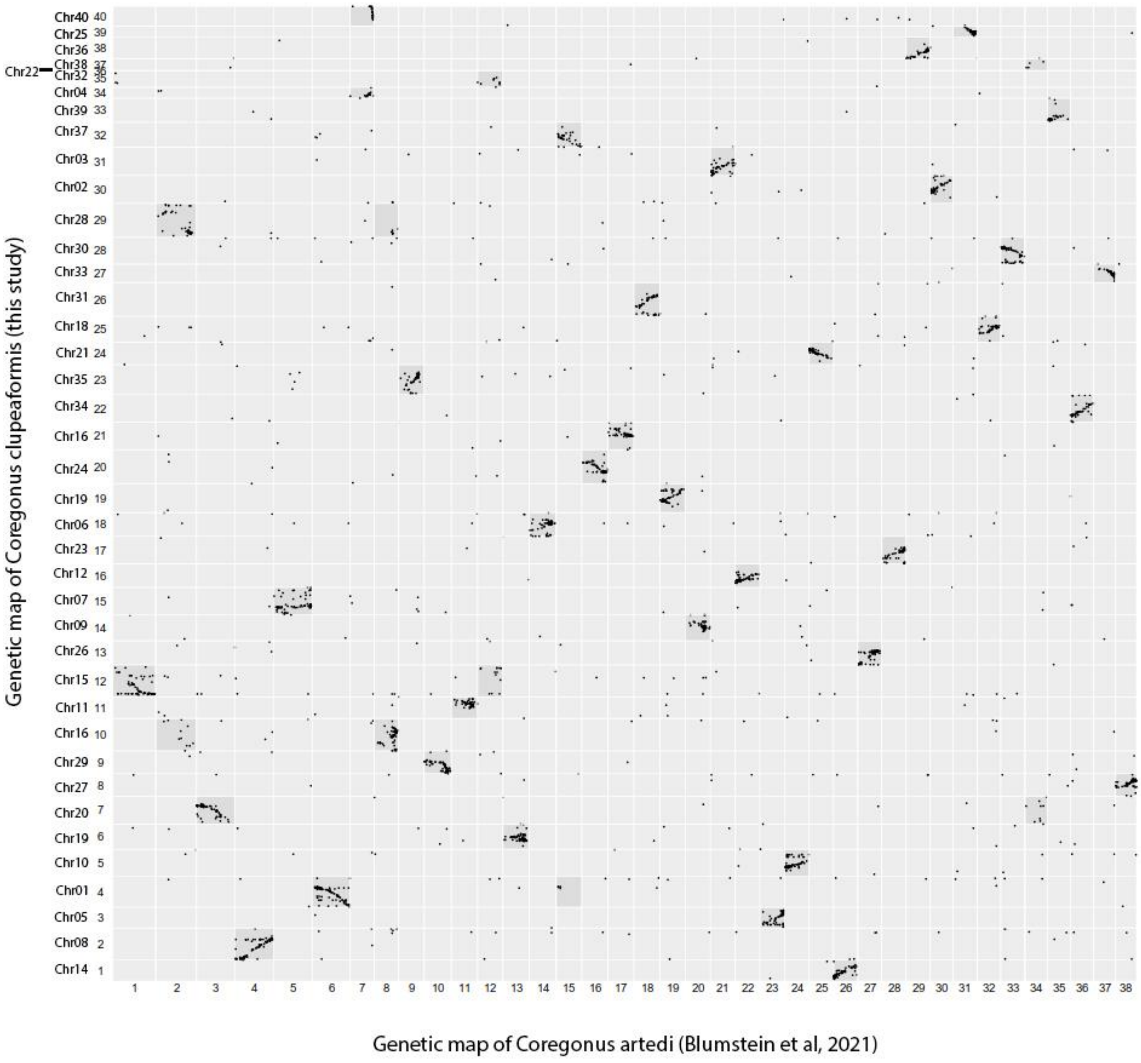

Figure S1: Homologous chromosomes with *Coregonus artedii*

*Coregonus clupeaformis* compared with *Coregonus artedii* (Blumstein et al., 2020) with markers paired through *Coregonus clupeaformis* genome identified homology between chromosome arms with MapComp (Sutherland et al., 2016).

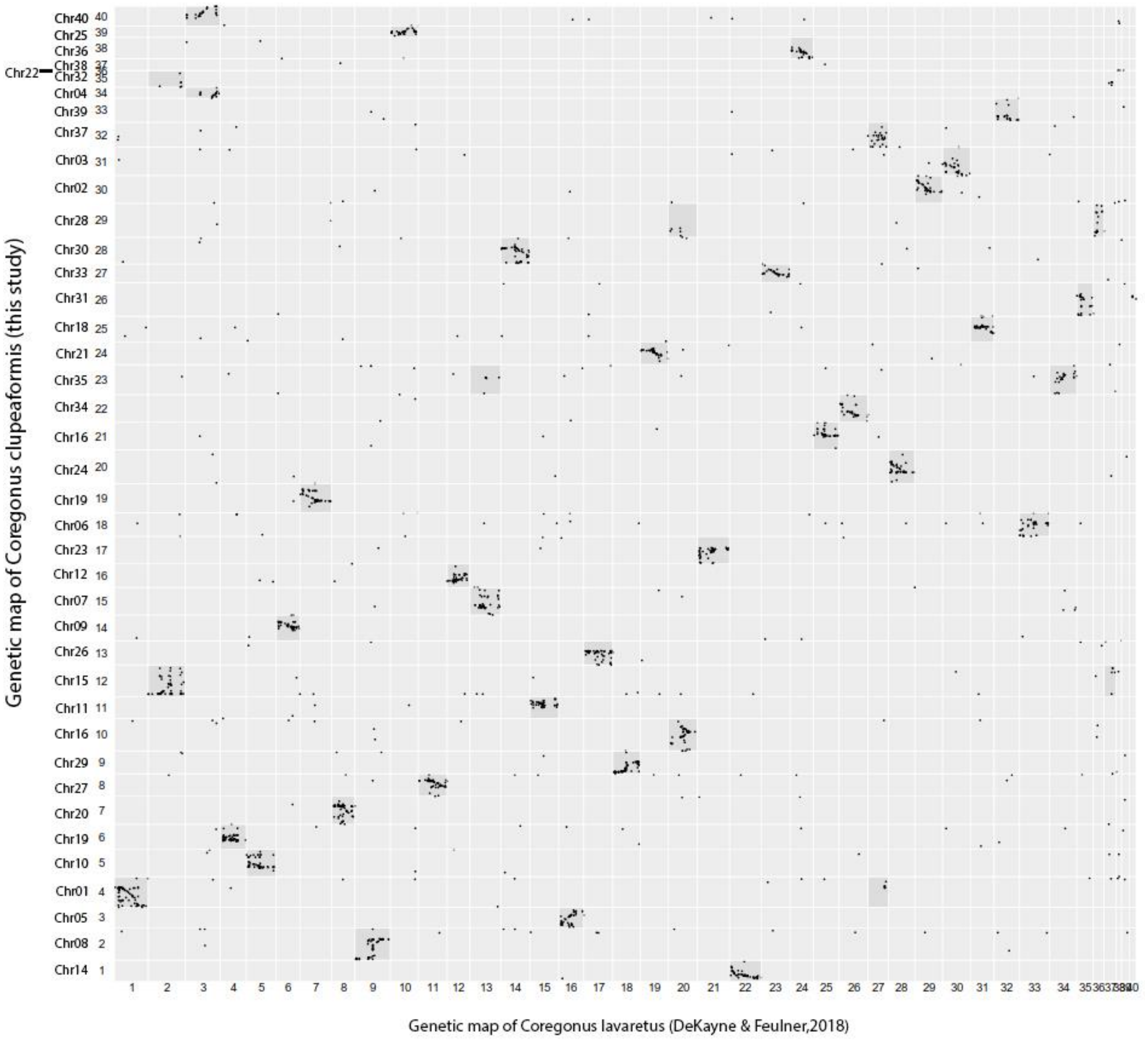

Figure S2: Homologous chromosomes with *Coregonus lavaretus*

*Coregonus clupeaformis* compared with *Coregonus lavaretus* (De-Kayne & Feulner, 2018) with markers paired through *Coregonus clupeaformis* genome identified homology between chromosome arms with MapComp (Sutherland et al., 2016).

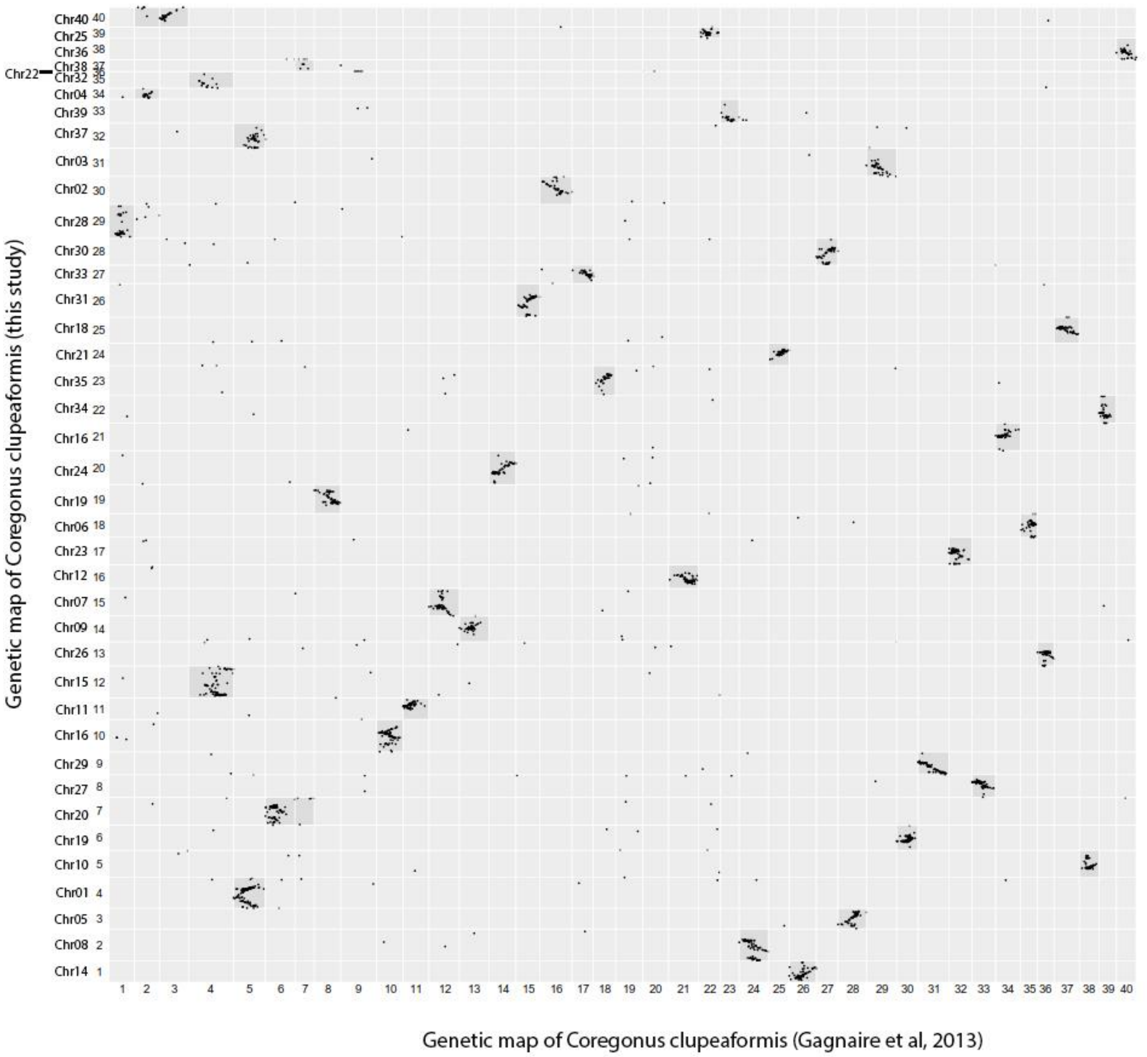

Figure S3: Homologous chromosomes with the previous map of *Coregonus clupeaformis*

*Coregonus clupeaformis* compared with *Coregonus clupeaformis* (Gagnaire et al., 2013) with markers paired through *Coregonus clupeaformis* genome identified homology between chromosome arms with MapComp (Sutherland et al., 2016).

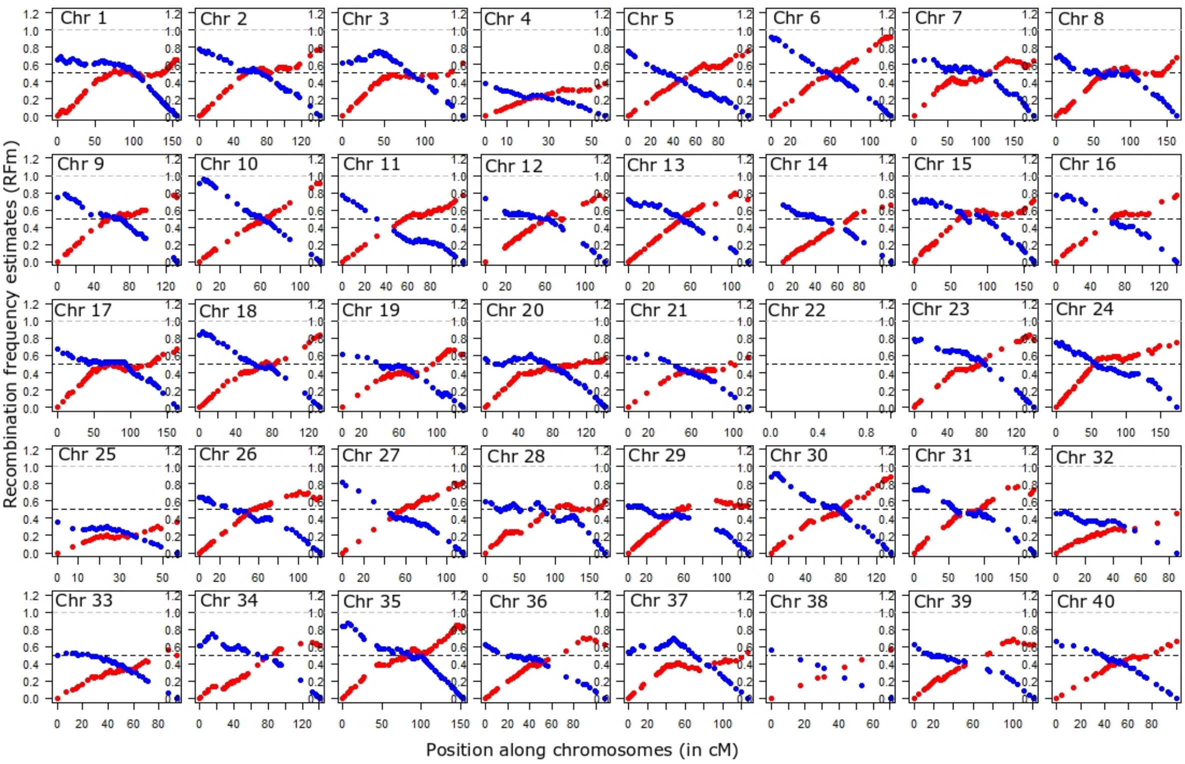

Figure S4: Recombination frequency estimates (RFm) for intervals between markers along each of the 40 linkage groups (LG).

LG are ordered by chromosome names from left to right, then from top to bottom). For each LG, RFm was calculated from both chromosomal extremities (right: red circles; left: blue circles), using each of the two terminal markers as a reference starting point. The RFm plot of Chr01 (top left) illustrates a classical metacentric pattern with a centromere position ~100cM while Chr05 (5th on the 1st line) illustrates a classical acrocentric pattern, the centromere position remains undetermined with regard to which LG extremity. See (Limborg, McKinney, Seeb, & Seeb, 2016) for methods.

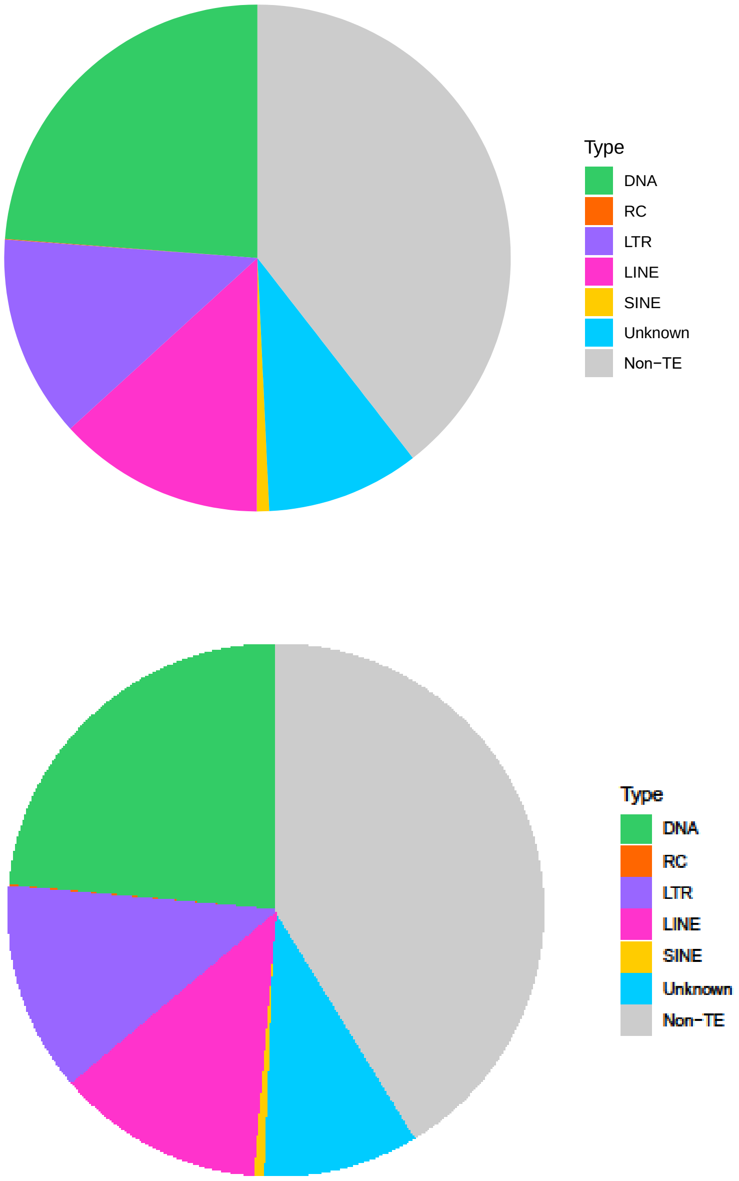

Figure S5: Proportion of transposable elements in interspersed repeats

Top: *Coregonus clupeaformis* sp. Normal (DNA=DNA-TIR 24%, RC-Helitron <1%, LTR 13%, LINEs 13%, SINEs <1%, Unknown=Unclassified TEs 9%, non-TE= interspersed repeats which are not transposable elements, 40%).

Bottom: *Coregonus clupeaformis* sp. Dwarf (DNA=DNA-TIR 23.5%, RC-Helitron <1%, LTR 13%, LINEs 13%, SINEs <1%, Unknown=Unclassified TEs 9%, non-TE= interspersed repeats which are not transposable elements, 40%).

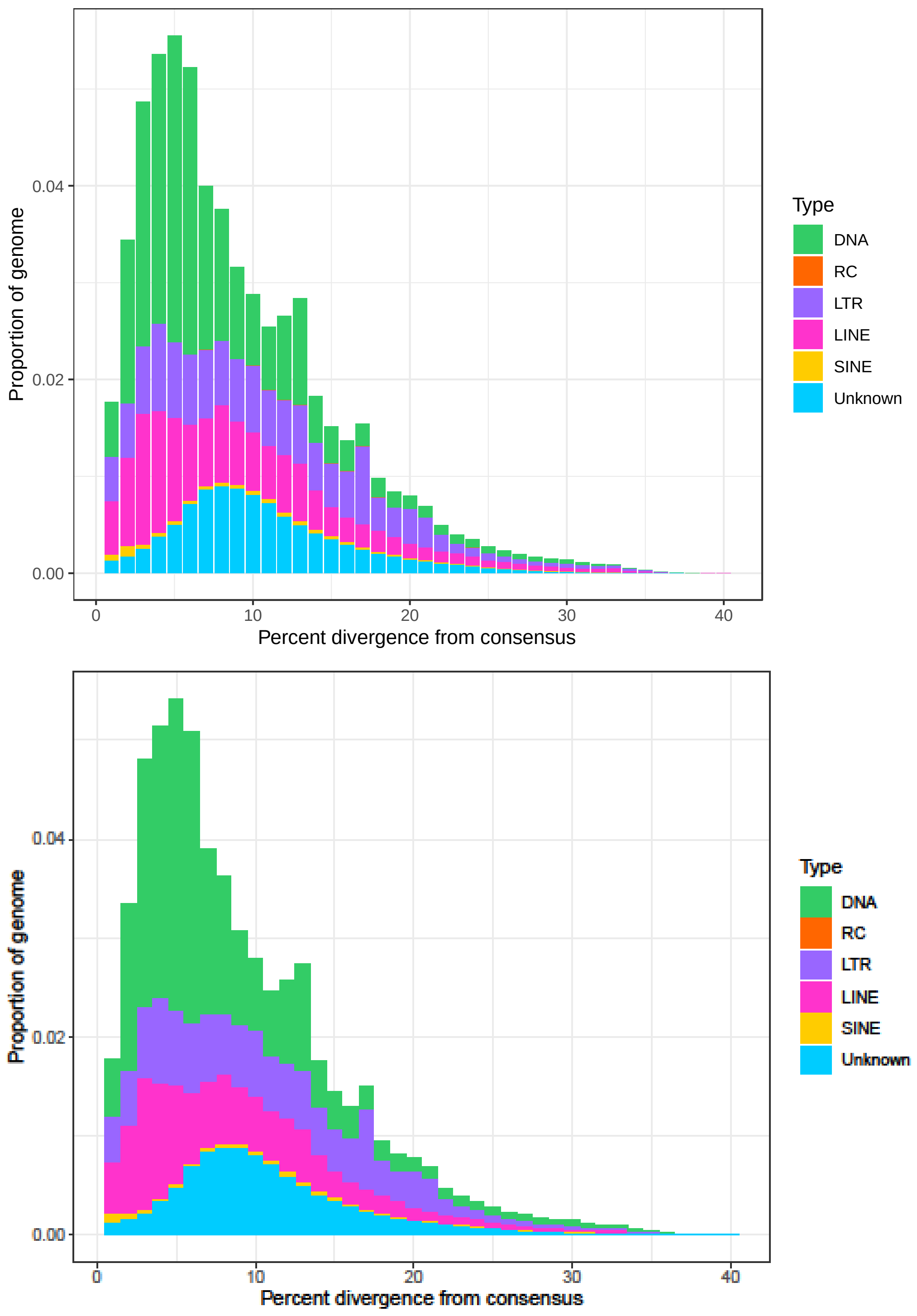

Figure S6: Distribution of transposable elements according to their divergence from the consensus.

Top: *Coregonus clupeaformis* sp. Normal. Bottom: *Coregonus clupeaformis* sp. Dwarf.

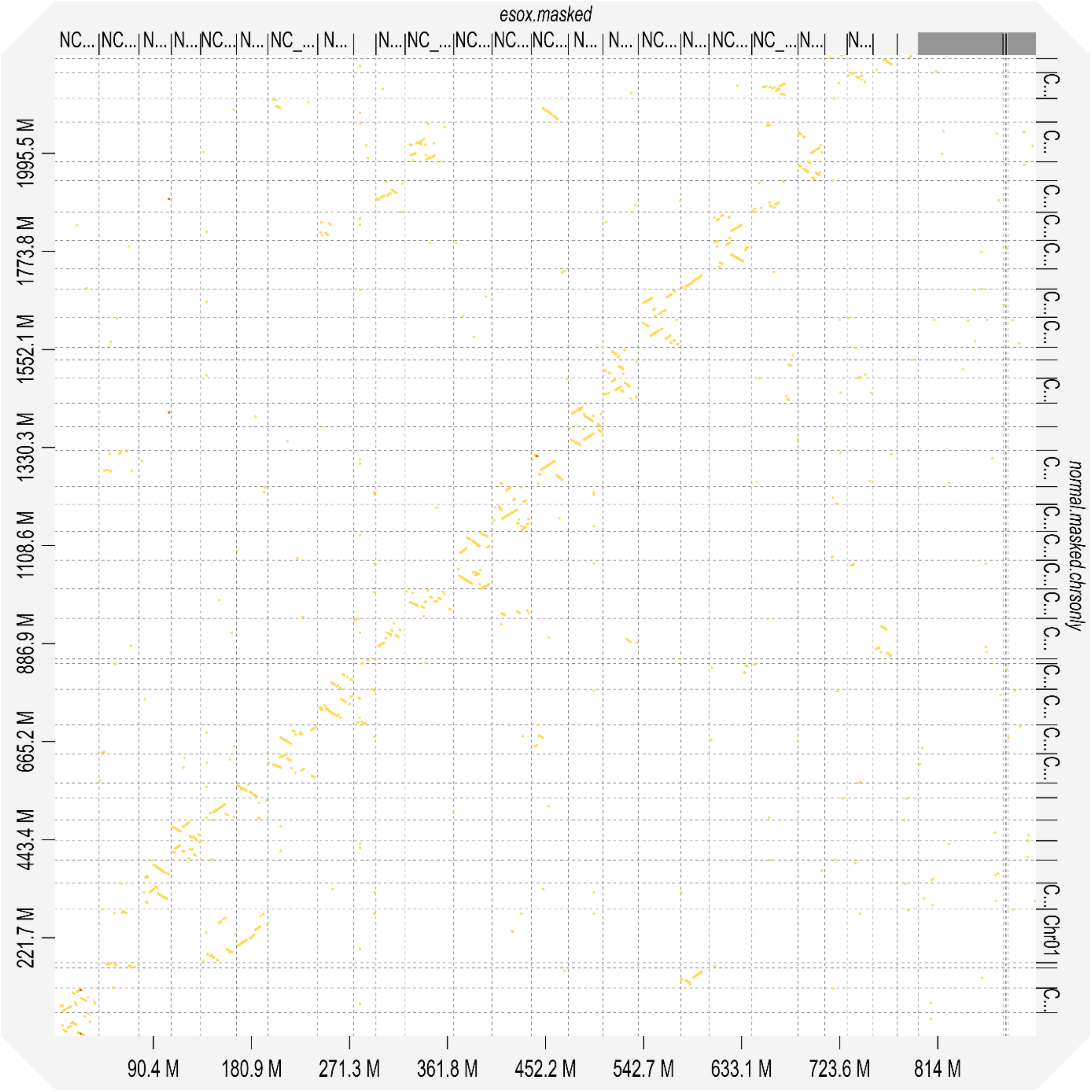

Figure S7: Alignment of the Normal Lake Whitefish genome to the Northern Pike genome using D-genies visualisation.

(Cabanettes & Klopp, 2018).

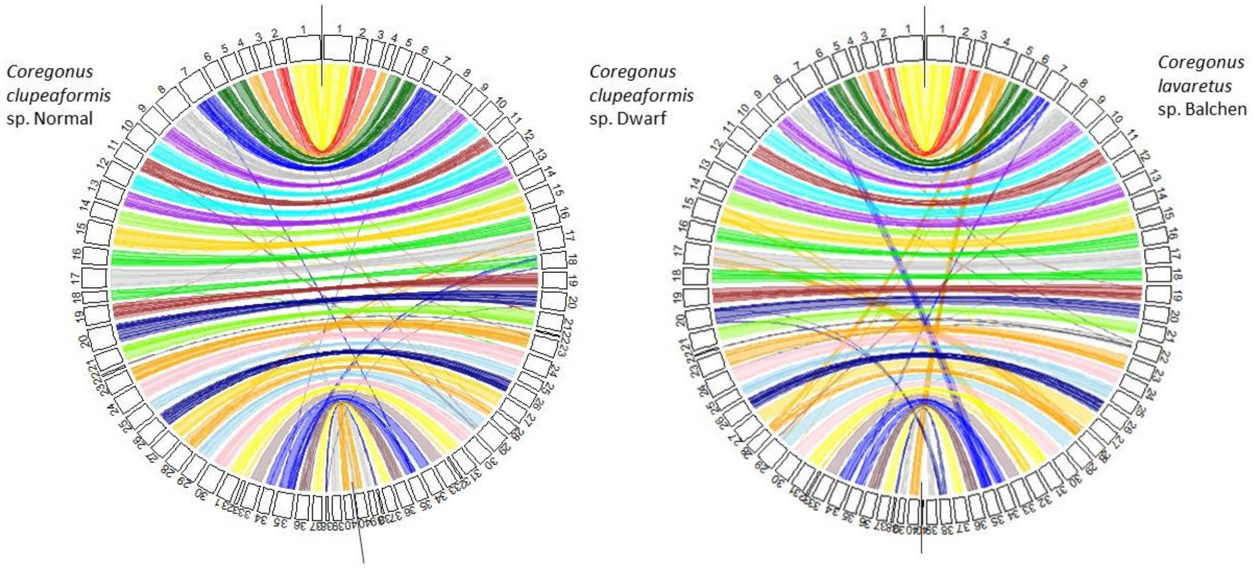

### Figure S8: Synteny between *C. clupeaformis* sp. Dwarf, *C. clupeaformis* sp. Normal, and *C. lavaretus* sp. Balchen

Circular plots showing syntenic relationship between *C. clupeaformis* sp. Dwarf and *C. clupeaformis* sp. Normal (left) and *C. lavaretus* sp. Balchen (right). Note that The Dwarf genome was anchored into chromosomes using the same linkage map as the Normal genome (based on a hybrid family).

### References in supplementary materials

Blumstein, D. M., Campbell, M. A., Hale, M. C., Sutherland, B. J., McKinney, G. J., Stott, W., & Larson, W. A. (2020). Comparative genomic analyses and a novel linkage map for cisco (Coregonus artedi) provide insights into chromosomal evolution and rediploidization across salmonids. *G3: Genes, Genomes, Genetics*, *10*(8), 2863–2878.

Cabanettes, F., & Klopp, C. (2018). D-GENIES: dot plot large genomes in an interactive, efficient and simple way. *PeerJ*, *6*, e4958.

De-Kayne, R., & Feulner, P. G. (2018). A European whitefish linkage map and its implications for understanding genome-wide synteny between salmonids following whole genome duplication. *G3: Genes, Genomes, Genetics*, *8*(12), 3745–3755.

Gagnaire, P., Normandeau, E., Pavey, S. A., & Bernatchez, L. (2013). Mapping phenotypic, expression and transmission ratio distortion QTL using RAD markers in the Lake Whitefish (Coregonus clupeaformis). *Molecular Ecology*, *22*(11), 3036–3048.

Limborg, M. T., McKinney, G. J., Seeb, L. W., & Seeb, J. E. (2016). Recombination patterns reveal information about centromere location on linkage maps. *Molecular Ecology Resources*, *16*(3), 655–661.

Rogers, S. M., Isabel, N., & Bernatchez, L. (2007). Linkage maps of the dwarf and normal lake whitefish (Coregonus clupeaformis) species complex and their hybrids reveal the genetic architecture of population divergence. *Genetics*, *175*(1), 375–398.

Sutherland, B. J. G., Gosselin, T., Normandeau, E., Lamothe, M., Isabel, N., Audet, C., & Bernatchez, L. (2016). Salmonid Chromosome Evolution as Revealed by a Novel Method for Comparing RADseq Linkage Maps. *Genome Biology and Evolution*, *8*(12), 3600–3617. doi: 10.1093/gbe/evw262
